# Targeted redesign of suramin analogs for novel antimicrobial lead development

**DOI:** 10.1101/2021.05.17.444489

**Authors:** Debayan Dey, Suryanarayanarao Ramakumar, Graeme L. Conn

## Abstract

The emergence of new viral infections and drug resistant bacteria urgently necessitates expedient therapeutic development. Repurposing and redesign of existing drugs against different targets is one potential way in which to accelerate this process. Suramin was initially developed as a successful anti-parasitic drug but has also shown promising antiviral and antibacterial activities. However, due to its high conformational flexibility and negative charge, suramin is considered quite promiscuous towards positively charged sites within nucleic acid binding proteins. Although some suramin analogs have been developed against specific targets, only limited structure activity relationship (SAR) studies were performed, and virtual screening has yet to be used to identify more specific inhibitor(s) based on its scaffold. Using available structures, we investigated suramin’s target diversity, confirming that suramin preferentially binds to protein pockets which are both positively charged and enriched in aromatic or leucine residues. Further, suramin’s high conformational flexibility allows adaptation to structurally diverse binding surfaces. From this platform, we developed a framework for structure- and docking-guided elaboration of suramin analog scaffolds using virtual screening of suramin and heparin analogs against a panel of diverse therapeutically relevant viral and bacterial protein targets. Use of this new framework to design potentially specific suramin analogs is exemplified using the SARS-CoV-2 RNA-dependent RNA polymerase (RdRp) and nucleocapsid protein, identifying leads that might inhibit a wide range of coronaviruses. The approach presented here establishes a computational framework for designing suramin analogs against different bacterial and viral targets and repurposing existing drugs for more specific inhibitory activity.

**For Table of Contents Use Only:** Table of Contents Graphic

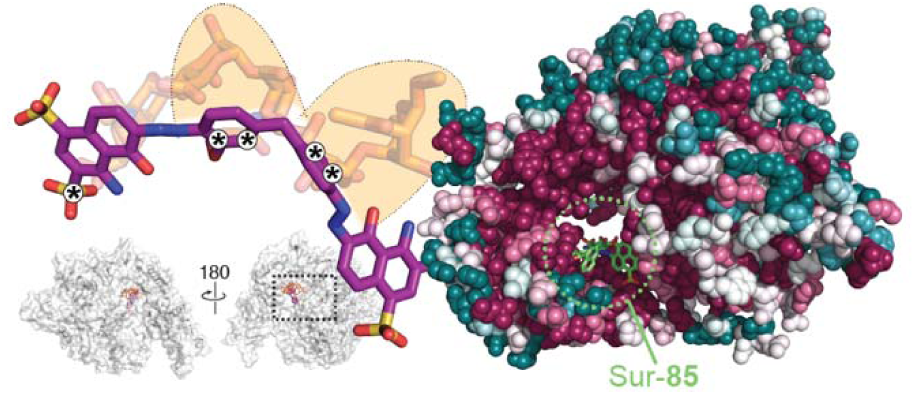

## INTRODUCTION

The rise of multi-drug resistant bacteria and emergence of new infectious viral pathogens, as exemplified by SARS-CoV-2 the cause of the COVID-19 pandemic, are dual major threats to human health world-wide. Effective new antimicrobials are thus urgently needed, but processes for their identification and development are often long and expensive. However, repurposing of existing, approved drugs and redesign of their activity against new targets represents one of the most promising opportunities to by-pass many of these hurdles^1^.

The unique challenges posed by COVID-19 have resulted in testing of a large array of antivirals, anti-inflammatory drugs, and other potential inhibitors of receptors of viral attachment for their effectiveness against SARS-CoV-2. For example, the nucleoside analog remdesivir, originally developed for Ebola virus, was found to be effective against SARS-CoV-2 with high selectivity, leading to its use for treatment of COVID-19 in many countries^2^. Other antivirals like favipiravir, ribavirin, darunavir, and lopinavir-ritonavir combination are either already in use or in clinical trials for SARS-CoV-2 treatment^3^. In addition, non-antivirals drugs like dexamethasone, which is used commonly to treat inflammatory and autoimmune conditions^4^, and the anti-parasitic ivermectin have shown potential as COVID-19 treatments^5, 6^.

Suramin was among the first anti-infective agents developed in the early 1920s and has also recently been shown to inhibit the progression of SARS-CoV-2 infection in human airway epithelial cell culture model^7^. Suramin appears on the World Health Organization’s List of Essential Medicines and has a long history of clinical use, with application over the last century in treating African Trypanosomiasis (sleeping sickness) caused by *Trypanosoma brucei*^8^, and other parasitic infections including onchocerciasis (African river blindness), leishmaniasis and malaria^9–11^. With its long history of successful use as an anti-parasitic, suramin and its analogs have also found use in wide variety of other treatments, such as anti-cancer agents, antivirals, and venom antidotes^12^.

Suramin was found to exhibit broad antiviral properties through distinct activities against components of diverse viruses. For example, through binding to glycoprotein gp120 suramin inhibits attachment of HIV to host T-cells^13^. Suramin also limits host cell entry of several other viruses, including herpes, dengue and hepatitis C virus^14–16^ and also inhibits enteroviral attachment to human host cells by binding to their nucleocapsid protein^17^. Suramin also inhibits some processes in viral replication, for example through binding to Zika virus NS2B/NS3 proteinase^18^ or to the RNA-dependent RNA polymerase (RdRp) of norovirus, chikungunya virus and SARS-CoV-2^19–21^. Suramin and related compounds thus hold promise as an initial lead for multiple targets, including viral RdRp and nucleocapsid, as well as future structure-activity relationship (SAR) studies or structure-guided design of suramin analogs as new therapeutics.

Although suramin and its analogs have been studied for many decades, including some SAR studies^22, 23^, many fundamental questions remain regarding the origin of their promiscuity, binding pocket preferences, and mode of on-target interaction. Suramin’s effectiveness in targeting viral nucleocapsid and RdRp has not yet been exploited to rationally design more selective and less toxic analogs based on this scaffold. Moreover, to our knowledge, no systematic high-throughput or virtual screening has been performed for suramin analogs for any viral targets (e.g. RdRp or nucleocapsid) to explore potential lead molecules. Here, we examine the structural basis of suramin’s ability to bind to diverse proteins and use these insights to develop a framework involving virtual screening, precision docking and docking-guided elaboration of the suramin scaffold against different therapeutically important bacterial and viral targets. Docking studies with a panel of nine diverse proteins reveals high-scoring suramin analogs to be more conformationally selective than suramin and to have greater target specificity. Finally, SARS-CoV-2 RNA-dependent RNA polymerase (RdRp) and nucleocapsid are used to exemplify the process of suramin analog redesign to generate novel analogs with the potential to bind with higher affinity and target specificity. Evolutionary analyses suggest that the optimized leads could potentially inhibit a wide range of coronaviruses^24^. This study thus sets out a computational framework using structure- and docking-guided ligand elaboration that can support redesign and repurposing existing antimicrobials for new, specific inhibitory activities.

## RESULTS

### Structural basis for suramin protein target site diversity

Suramin is a C2 symmetric molecule with each half comprised of a trisulfonated naphthyl ring with negative charge at physiological pH, connected via amide linkages to toluene and phenyl rings (**Fig. 1A**). Suramin is highly flexible with a total of 10 rotatable bonds (**Fig. 1A**; α_1-5_ and α_1’-5’_) and no substituent groups on the four central rings that might otherwise restrict its conformational freedom. This flexibility, and thus conformational adaptability, is likely a key factor in suramin’s ability to interact with diverse target sites in many proteins, but also its high off-target binding. Similarly, another polyanionic class of molecules similar to suramin, the glycosaminoglycans (GAGs) such as heparin (**Fig. 1B**), can bind multiple diverse protein targets but also suffer the limitation of high off-target binding.

**Figure 1.**
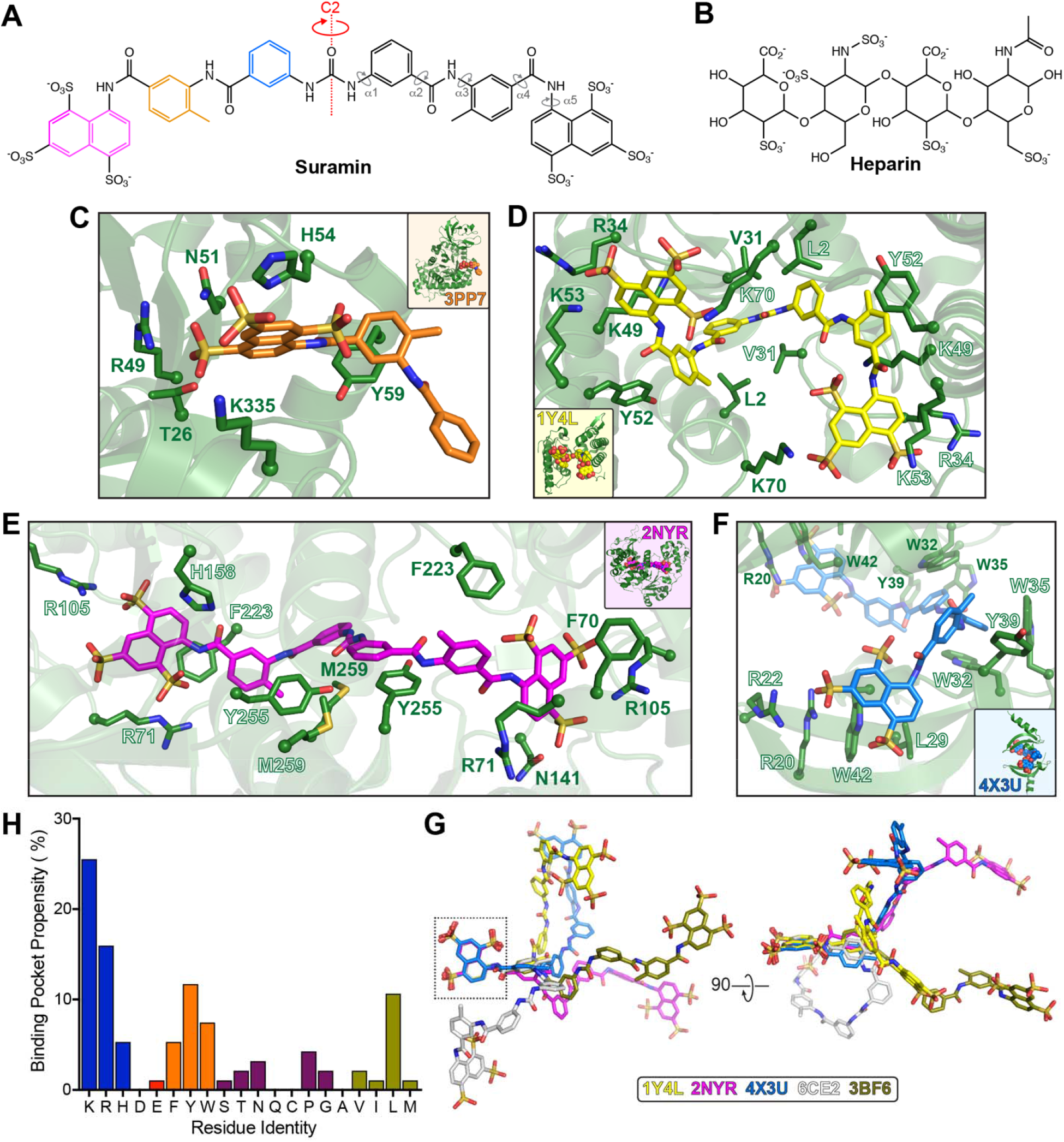
Structural basis for suramin protein target site diversity. Chemical structures of ***A***, suramin and ***B***, heparin. Crystal structures of suramin (or partial fragment) bound to: ***C***, Leishmania mexicana pyruvate kinase (PDB 3PP7), ***D***, snake venom protein myotoxin (PDB 1Y4L), ***E***, NAD^+^-dependent protein deacetylase SIRT5 (PDB 2NYR), and ***F***, epigenetic reader chromobox homolog 7 (PDB 4X3U). ***H***, Residue propensities in the suramin binding pockets in targets shown in *panels C-F* and **Figure S1**. ***G***, Superposition via the boxed sulfo-naphthyl ring of suramin molecules from co-crystal structures with diverse targets (PDB 1Y4L, 2NYR, 4X3U, 6CE2 and 3BF6).

To begin defining the residue preferences in suramin binding pockets, we examined structures of several suramin complexes with diverse proteins (**Table S1**). These structures included the *Leishmania mexicana* pyruvate kinase (PDB 3PP7**; Fig. 1C**) with suramin bound in its ATP binding site, snake venom protein myotoxin (PDB 1Y4L; **Fig. 1D**), and two human protein targets for treatment of cancer, metabolic and neurological diseases, the NAD^+^-dependent protein deacetylase sirtuin 5 (SIRT5; PDB 2NYR; **Fig. 1E**) and the epigenetic reader chromobox homolog 7 (PDB 4X3U; **Fig. 1F**)^10, 25–27^. A number of general features of the interaction networks within the suramin binding pockets are immediately apparent. The trisulfo-naphthyl ring interacts with positively charged residues (Arg, Lys, His) via its sulfate groups while the naphthyl ring makes π-mediated interactions with aromatic residues, Phe, His, Tyr and Trp (**Fig. 1C-F**). Aromatic residues with a polar ring atom may also make a hydrogen bonding interaction with a sulfonate group, coordinating both features of the trisufonated naphthyl ring. The toluene and central phenyl rings of suramin are additionally stabilized by networks of aromatic and hydrophobic residues (**Fig. 1C-F**).

Suramin complexes with viral proteins have also been determined for norovirus RdRp (PDB 3UR0)^19^ and bunyavirus nucleocapsid (PDB 4J4V)^28^, but these offer only partial snapshots of favored suramin binding sites as just a portion of suramin could be modeled in each crystal structure (**Fig. S1**). In the RNA binding channel of norovirus RdRp34, the trisulfo-naphthyl ring of suramin is anchored by both positive (Lys171 and Arg392) and aromatic (Trp42) residues, while the central region is stabilized by Gln66, Lys180, Lys181, Arg245 and Lys68. The bunyavirus nucleocapsid forms a pentameric complex with an RNA binding cavity on the inner edge of the ring-like assembly where suramin also interacts^28^. Again, each of the modeled suramin rings is stabilized by multiple ionic, hydrogen bonding and aromatic/ hydrophobic stacking interactions, including residues Asn66, Lys67 and Arg95 with the trisulfo-naphthyl ring, and Arg64, Thr63, Gly65, Phe177 and Pro127 the two central rings (**Fig. S1B**). Thus, visual inspection of these viral protein structures reveals overall consistent trends in suramin’s binding pocket residue interaction preferences with other proteins of diverse origin and function.

More detailed analyses of available suramin-bound protein structures reveal consistent features in the binding pocket physicochemical composition, including strong enrichment of specific residues (**Fig. 1G**). Positively charged Lys and Arg are highly enriched in the pocket compared to the full protein (25% and 15%, respectively, of all interacting residues), while His is more modestly enriched (5%), but, as noted, can make both aromatic stacking and charged interactions. The absence of negatively charged residues (~1.5% combined) also points towards suramin’s strong preference for positively charged binding regions, such as nucleic acid binding sites. In total, polar residues are slightly less favored (~12 % total) as compared to aliphatic hydrophobic residues (~14%), with a marked preference for Leu (10.6 %) among the latter group (**Fig. 1G**). This may be due to leucine’s aliphatic stacking potential with the aromatic rings of suramin. Aromatic residues (Trp, Tyr and Phe; together ~25% of total) are also strongly enriched in suramin binding sites.

Suramin’s binding promiscuity for diverse protein targets is likely underpinned by its high inherent conformational flexibility, as revealed by superposition of protein-bound suramin molecules using one trisulfo-naphthyl ring (**Fig. 1H**). Suramin bound to thrombin (PDB 3BF6) adopts the most extended conformation we identified (and is used hereafter as the reference), with the trisulfo-naphthyl rings ~29 Å apart. Suramin bound to SIRT5 (PDB 2NYR) also has a linear conformation but with differences in the α2, α3, α2’, and α3’ angles (i.e. the dihedrals between the toluene and phenyl rings). In contrast, suramin bound to myotoxin I (PDB 6CE2) has its trisulfo-naphthyl rings only ~14 Å apart and is observed in a highly bent conformation. Similarly, suramin bound to myotoxin II (PDB 1Y4L) adopts a highly bent conformation, but distinguished by additional changes in the α1 and α1’ dihedral angles. To further assess the low energy conformations of suramin and two analogs, NF449 and NF023, we used a computational *Monte Carlo* conformational scan which revealed 116 potential low energy conformational states for suramin, while NF449 and NF023 each have 143 and 80, respectively.

Collectively, these structural and computational conformation analyses confirm suramin to be a highly flexible ligand, with multiple potential hinge points, that can adopt a large variety of conformations to fit a given target protein binding pocket. Further, these favored binding pockets are positively charged and enriched with aromatic and Leu residues, such as most commonly occur in large interfaces or in protein regions which bind to nucleic acids or nucleotides.

### Identification of suramin and heparin analogs targeting viral and bacterial proteins

Previous studies have identified inhibitory activity of suramin against bacterial and viral proteins, but no computational or high-throughput screens have been conducted with suramin analogs against such targets. Structure-guided analysis of suramin analogs bound to various viral or bacterial targets using large-scale docking studies could, however, allow for downstream rational design of novel analogs with improved affinity and specificity towards a desired target. We devised a strategy to accomplish this goal using docking studies with suramin and heparin analogs and using docking poses of the latter set of ligands to guide design of new suramin analogs with substituents positioned to make additional protein target-specific interactions (**Fig. 2**).

**Figure 2.**
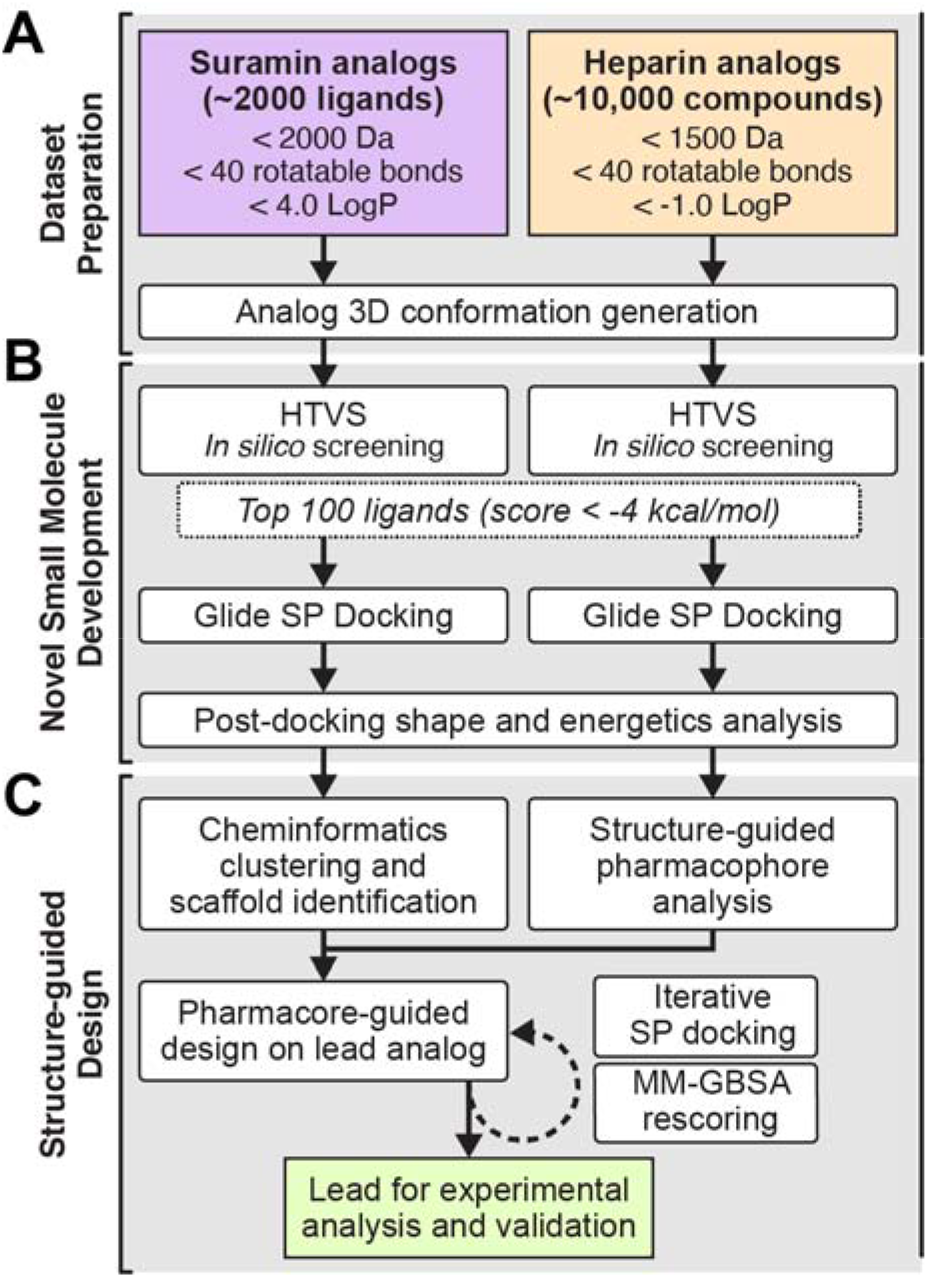
Computational workflow used in this study. ***A***, Suramin and heparin analog sets were retrieved from PubChem with the indicated parameter filters and subject to conformer generation in LigPrep (Schrödinger Software). ***B***, Suramin and heparin analog sets were used for separate HTVS and the resulting top 100 ligands then subject to higher precision (SP) docking in Glide (Schrödinger Software). ***C***, Following further shape and energetic analysis, chemoinformatics was used to determine different fragments of top scoring suramin analogs. For heparin analogs, the top poses corresponding to the top selected suramin analog for each target was next used for pharmacophore-based identification of regions in the suramin analog which could be substituted to improve its predicted affinity and target specificity. The iterative process of ligand re-design employed further SP docking and/or MM-GBSA. Identified novel analogs would next be used as leads for experimental analysis and validation.

Nucleic acid binding proteins in bacteria and viruses play crucial roles in the processing of genetic information and thus represent important targets for therapeutic development. Due to suramin’s affinity for nucleic acid binding proteins, a test panel of diverse protein targets known to interact with DNA or RNA was selected. Viral proteins, known to be bound by suramin, included RdRp of Norovirus and SARS-CoV-2, which is involved in replication and transcription of the viral RNA genome^19, 21^; nucleocapsid of bunyavirus and SARS-CoV-2, which is involved in packaging the viral RNA genome and virion assembly, and thus of critical importance in viral infection^17, 28, 29^; and, human immunodeficiency virus type 1 (HIV-1) protein gp120, for which binding to galactosylceramide on human colon epithelial cells is blocked by suramin^13, 30^. To further diversify the protein target set, we included four bacterial proteins: DNA-binding protein HU (HU), DNA recombination protein RecA (RecA), DNA-binding protein Fis (Fis), and the thiostrepton-resistance methyltransferase (TsnR). HU and Fis are global regulators which play important roles in bacterial gene regulation, biofilm development and maintaining nucleoid architecture^31–34^, while RecA is an essential DNA recombination and repair protein known to be targeted by suramin^35^. Finally, TsnR is a ribosomal RNA methyltransferase that confers resistance to the antibiotic thiostrepton^36^, and represents a wider group of RNA modifying enzymes that are promising targets for novel drugs to break antibiotic resistance. To generate the ligand sets, we next used a small panel of known biosimilars (**Table S2**) in a PubChem search which identified ~2,000 and ~10,000 analogs of suramin and heparin, respectively (**Fig. 2A**).

Using the protein target panel and suramin/ heparin analog sets, high-throughput virtual screening (HTVS; **Fig. 2B**) in the Glide module of the Schrödinger software produced a wide range of binding scores, typically ranging from −1.0 to −9.0 kcal/mol, but with large differences in distribution and average score for each protein target (**Fig. 3A,B**). Among the bacterial proteins, suramin analogs bind on average more tightly to Fis and TsnR (mean binding score −5.7 and −5.9 kcal/mol, respectively) compared to HU and RecA (−5.1 and −4.6 kcal/mol, respectively; **Fig 3A**). HIV protein gp120 was predicted to be bound less tightly with a mean score of −3.6 kcal/mol, whereas both suramin and heparin analogs docked viral RdRp with comparatively higher score (**Fig. 3A,B**).

**Figure 3.**
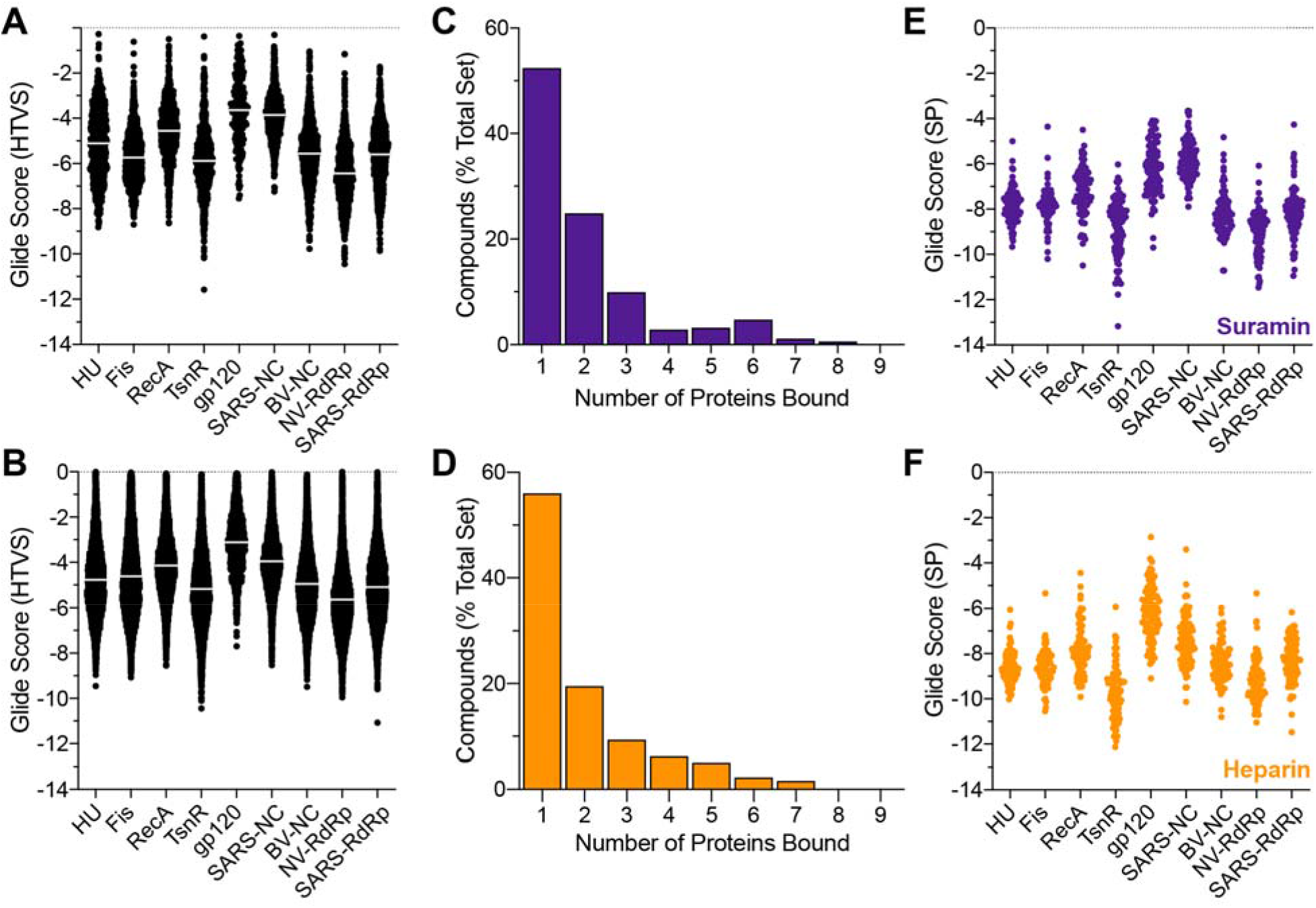
Virtual screening and precision docking identify suramin analogs with increased target specificity. HTVS docking scores for ***A,*** suramin and ***B,*** heparin analogs against bacterial targets HU, Fis, RecA and TsnR; and, viral targets gp120, SARS-CoV-2 and norovirus RdRp, and SARS-CoV-2 and bunyavirus nucleocapsid (SARS-NC and BV-NC, respectively). A high percentage of both ***C***, suramin and ***D***, heparin analogs bind to only one or two targets within the protein panel used, suggesting that virtual screening has identified more specific leads as compared to the parent molecules, suramin and heparin. ***E***,***F***, SP Glide docking scores for the top 100 ligands from HTVS for suramin and heparin analogs, respectively.

Among the top 100 scoring suramin and heparin analogs identified by HTVS, more than ~50% of these ligands bind to only a single protein among the nine targets (**Fig. 3C,D**). Approximately 30% of the remaining ligands in each set bind to only two or three receptors, with just a small fraction showing greater promiscuity (high docking score for 4 or more targets). This result indicates that virtual screening-based using suramin or heparin analogs can identify ligands with greater selectivity for diverse protein targets compared to using either suramin or heparin individually.

The top 100 suramin and heparin analogs from HTVS against each target protein were next subject to more accurate docking pose and binding score calculation using Standard Precision (SP) Glide (**Fig. 2B** and **Fig. 3E,F**). An improvement in the docking score was observed for many analogs due to the ligand conformational sampling used in the SP Glide docking method as compared to “rigid” ligand docking of HTVS. However, SP Glide docking had a very similar trend in docking score distribution for each set of top 100 analogs (**Fig. 3E,F**). Heparin analogs bind modestly better than those of suramin, likely due to their greater overall conformational flexibility and number of hydrogen bond donors/acceptors, which allow them more readily adapt to the wide variety of positively charged surfaces in these target proteins. The top scoring ligands from SP docking (**Tables S3** and **S4**, and Supplementary File “Top 100 docking results”) were then selected for subsequent elaboration of the suramin scaffold and are hereafter named beginning with “Sur” and “Hep” for suramin and heparin analogs, respectively.

### Strategy for docking-guided elaboration of the suramin scaffold

To facilitate suramin analog lead redesign for increased target selectivity and affinity, the top scoring suramin analogs for each protein target were visually inspected post-docking and then superposed with the top scoring heparin analog identified in the same binding site. The heparin analog set contains ligands that are typically more conformationally flexible and enriched in potential hydrogen bond donors/ acceptors. While these features would make the heparin analogs themselves poor leads for specific interactors, comparison of the bound analogs can direct identification of sites within the corresponding suramin analog that can be further substituted to make additional interactions with the target protein (**Fig. 2C**). As an example, structural analysis of the top suramin and heparin analogs directed against HU and the resulting suramin analog redesign strategy is described below.

HU interacts with DNA as a dimer via a positively charged binding cleft, lined with Lys and Arg residues from both protomers. Sur-1 binds to HU in this cleft with the highest affinity of the suramin analog set (−9.66 kcal/mol), placing two of its naphthyl rings over a surface formed by Pro81 and the adjacent β-strand of both HU protomers at the center of the dimer interface (**Fig. 4A**). Sur-1 is composed of three sulfo-naphthyl rings (**Fig. 4B**), with both terminal rings substituted with two sulfonyl groups, and a single sulfonate on the central ring. The majority of these groups are coordinated by multiple Arg residues, including Arg53, Arg55 and Arg80 from both protomers, and Arg58 and Lys 86 from a single protomer (**Fig. 4A**). Lys86 of the second protomer is also positioned near (~4-6 Å) the carbonyl and sulfonyl groups of the central and terminal naphthyl rings, respectively, exemplifying how the dimeric interface may be exploited by a pseudo-symmetrical suramin analog. Additionally, Sur-1 and other top HU-binding suramin analogs (Sur-2 and Sur-3; **Table S3**) are all markedly more conformationally rigid than suramin, due to the diazene linker between the naphthyl rings. Sur-1 has only four rotatable bonds with a single low energy conformation (**Fig. 4B**), compared to 10 rotatable bonds and 116 low energy conformations in suramin. The ligand docking process thus identified suramin analogs with both improved binding affinity to HU (based on docking score) and reduced conformational rigidity and net charge. These features make Sur-1 a superior candidate compared to suramin for developing a more specific inhibitor of HU.

**Figure 4.**
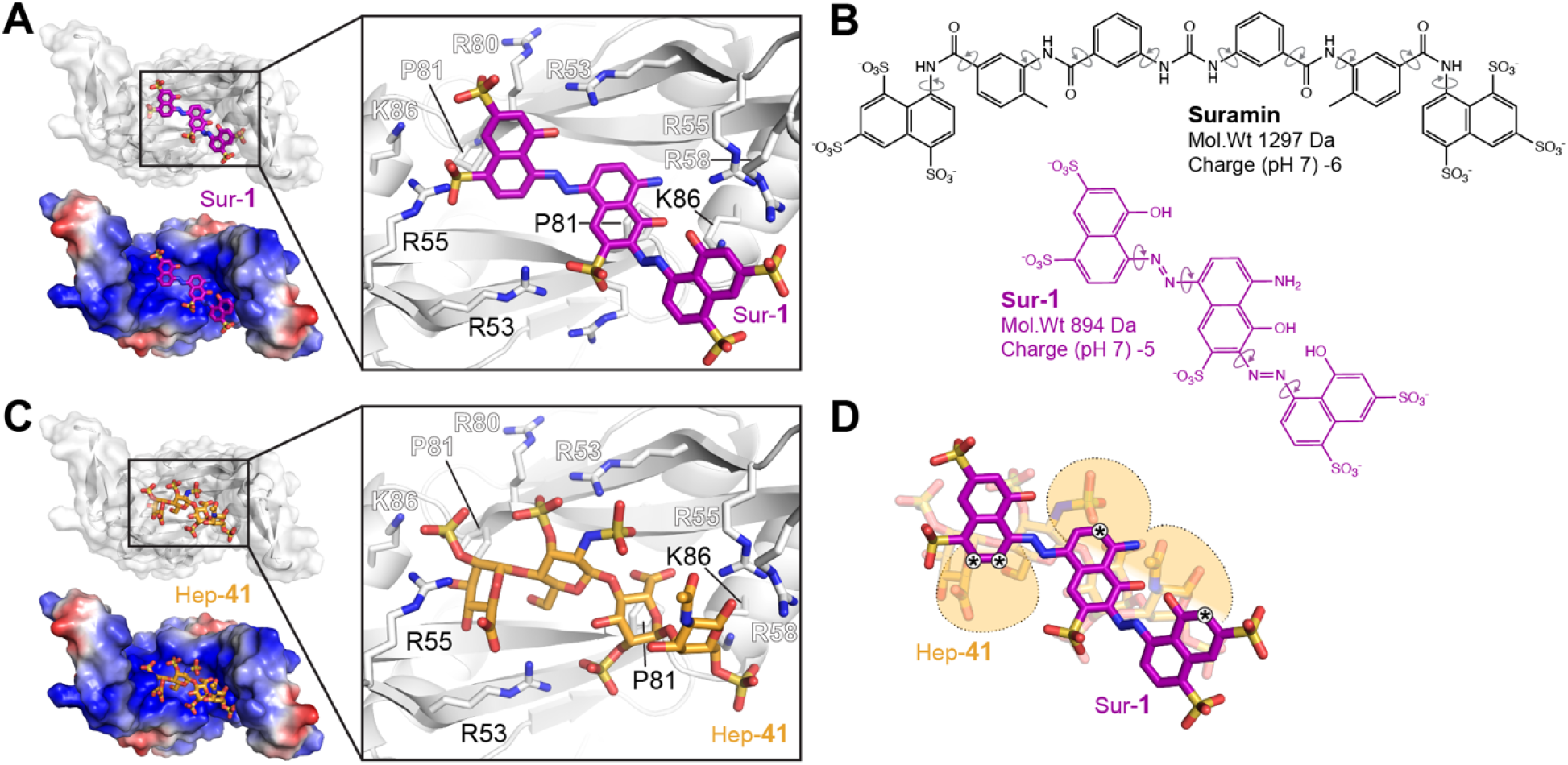
Docking-guided elaboration of the top identified suramin analog against HU. ***A,*** SP Glide docking pose of suramin analog Sur-1 (purple) bound to the DNA interaction surface of HU (top, white surface; bottom, electrostatic potential surface). ***B***, Comparison of the chemical structures of suramin and Sur-1, highlighting the reduced number of rotatable bonds in the analog. ***C,*** SP Glide docking pose of heparin analog Hep-1 (orange) bound to the same surface of HU. ***D***, The overlapping binding poses of Hep-1 and Sur-1 which were used to identify sites in Sur-1 for redesign (indicated *) to improve binding affinity and specificity.

Our docking procedure with the heparin analog set also identified Hep-41 as having high affinity (−10.02□kcal/mol) for the same pocket as Sur-1 in the DNA binding cleft of HU **(Fig. 4C**). Hep-41 is composed of sulfated iduronic acid and glucosamine with trioxidanylsulfanyl substitution, which make extensive interactions with essentially the same group of residues in the DNA binding cleft (i.e. Arg53, Arg55 Arg58, Arg80 and Lys86; **Fig. 4B**). However, while docking of Sur-1 and Hep-41 are qualitatively similar, due to its larger number of hydrogen bond donors/ acceptors, docking of the latter ligand offers detailed structure-guided cues for placement of new substituents in Sur-1. Specifically, superposition of the two analogs reveals sites on Sur-1 for optimal geometric placement of additional polar groups to enhance interaction within the binding site (**Fig. 4A,D**). For example, addition of polar groups on each naphthyl ring could increase target-ligand interactions by further engaging Arg53 and Arg55 (from both protomers) via bifurcated hydrogen bonding. These dual ligand docking comparisons can thus be used as a guide to improve the design of suramin analogs to increase both their target specificity and binding affinity.

This same strategy was used for the other eight target proteins, with heparin analog superposition identifying multiple sites in each top suramin analog where substitutions could be made to improve affinity and target selectivity (marked * in **Fig. S2-S5**; also see Supplemental Results for further details). Specifically, we identified sites for targeted redesign of Sur-6 (using Hep-42) for binding to the positively charged cleft of TsnR in an extended conformation (**Fig. S2 A**-**D**), and Sur-13 (using Hep-45) and Sur-15 (using Hep-50) for binding with higher affinity and specificity to Fis (**Fig. S2 E-H**) and RecA (**Fig. S3 A-D**), respectively. Among the viral proteins, HIV gp120 (6IEQ) binds to smaller analogs than the other targets with the highest docking score to Sur-22 and sites for targeted redesign identified by docking of Hep-51 (**Fig. S3E-H**). Similarly, for norovirus RdRp **(Fig. S4A**-**D**) and bunyavirus nucleocapsid **(Fig. S4E, F**) the docking identified Sur-24/ Hep-56 and Sur-27/ Hep-52 binding poses, respectively, for targeted redesign. Corresponding results for SARS-CoV-2 RdRp and nucleocapsid are described in the next section along with subsequent redesign and further docking analyses. In summary, virtual screening and precision docking analyses using suramin and heparin analog chemical space has discovered new lead analogs and a path to rational redesign for specific viral and bacterial protein targets.

### Suramin analog redesign for specific, high-affinity lead inhibitors of SARS-CoV-2 RdRp and nucleocapsid

SARS-CoV-2 RdRp is a large macromolecular complex composed of nsp12 catalytic subunit, with an RNA binding tunnel that presents a potential site for drug targeting (**Fig. S5A,B**), and nsp7-nsp8 cofactors^37^. Virtual screening and docking analyses identified Sur-36 as binding with highest docking score to SARS-CoV-2 RdRp, with Hep-58 also binding in the same region (**Fig. S5C**). Sur-36 has a central biphenyl ring attached to terminal sulfo-naphthyl groups joined by a conformationally restricted diazene linker (**Fig. 5B**). Two other similarly high-scoring ligands, Sur-37 and Sur-38, were also identified (**Table S3**), but Sur-36 was selected as the lead candidate due to the greater flexibility of these additional hits. Inspection of the RdRp binding pocket reveals that each ring of Sur-36 is stabilized by multiple interactions: the distal sulfo-naphthyl ring with Lys551, Ser549 and Arg553, the central phenyl rings with Ile548 and Ala549, and the proximal sulfo-naphthyl ring with Tyr546 and Asn497 (**Fig. 5A, B**).

**Figure 5.**
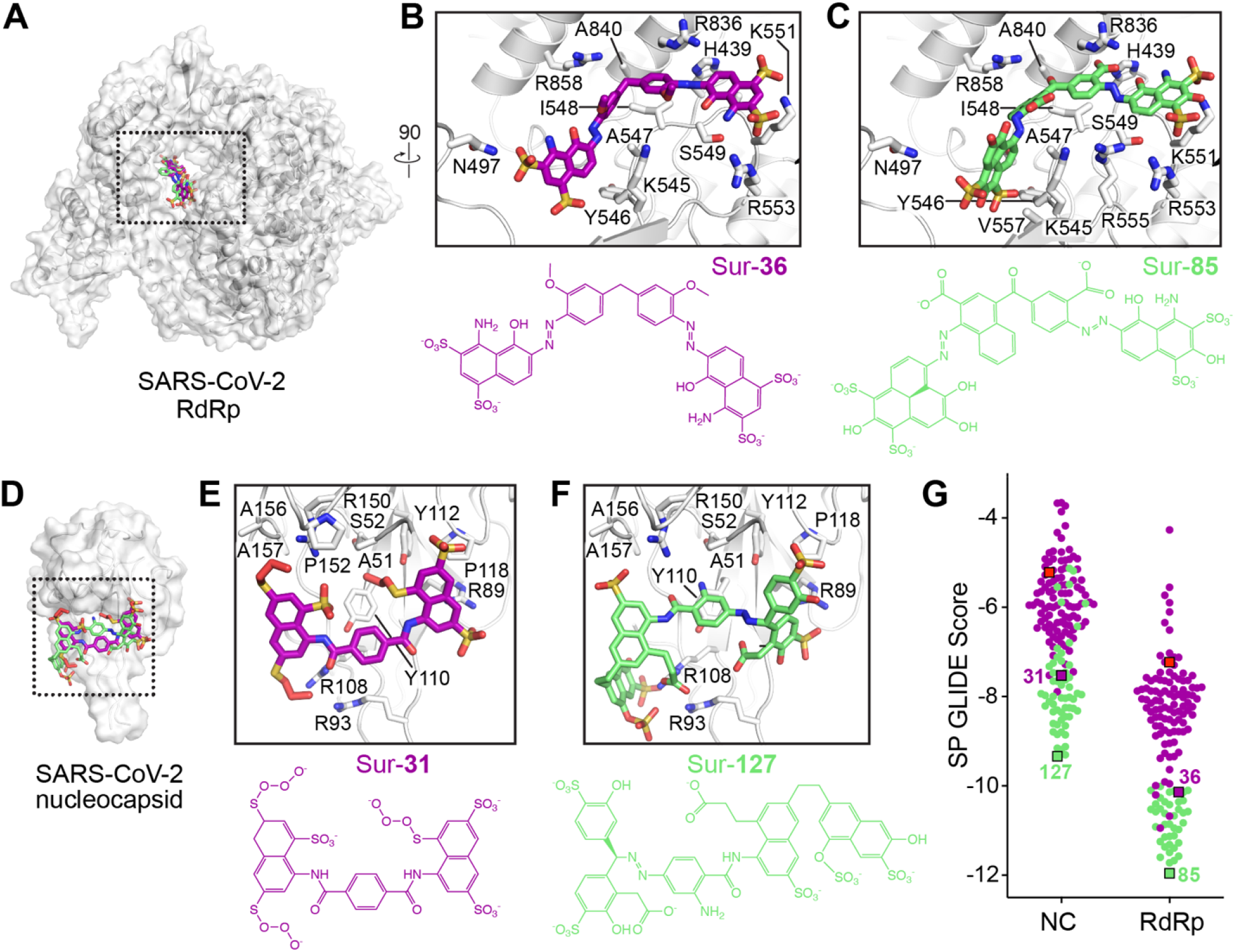
Design of suramin analogs specific to SARS-CoV-2 RdRp and nucleocapsid. ***A***, Location of suramin analogs Sur-36 and Sur-85 in the RNA binding tunnel of SARS-CoV-2 RdRp. ***B***, Zoomed in views of the boxed region in *panel A* of the SP Glide docking pose of Sur-36, an existing analog of suramin, bound to the RNA binding tunnel. ***C***, Iterative rounds of chemical enumeration (**Figure S6**) and docking, resulted in the designed suramin analog Sur-85shown in the same binding pocket. ***D***, Location of suramin analogs Sur-31 and Sur-127 on the RNA binding surface of SARS-CoV-2 nucleocapsid. ***E***,***F***, Zoomed in views of the boxed region in *panel D* of the SP Glide docking poses of Sur-31 and Sur-127, respectively. ***G***, Comparison of SP Glide docking scores for the initial top 100 suramin analogs (purple) and analogs tested during the iterative chemical enumeration process (green), showing the improvement in predicted affinity. Suramin (red square), starting analogs (Sur-31 and Sur-36 for nucleocapsid (NC) and RdRp, respectively) and final redesigned analogs (Sur-127 and Sur-85) are highlighted.

Using the Hep-58 binding pose, four positions for additional substitutions in Sur-36 were identified (**Fig. S5D**), as described above for the other ligand/ protein target pairs. A process of chemical enumeration, structure-guided design and iterative docking was then used to systematically alter the functional groups of substituents at the identified attachment positions and calculate the changes in the docking score (**Table S5**). The methoxy groups in the central phenyl rings were first modified to carboxylate, phenyl-methanol, halides and other functional groups, with docking analysis identifying the carboxylate groups (Sur-60; **Fig. S6A**) as conferring the greatest increase in docking score, from −10.14 (Sur-36) to −11.24 kcal/mol (Sur-60). In Sur-60, one phenyl ring carboxyl group engages Lys545, while the second carboxylate makes new contacts with Arg555. Additional docking-guided modifications were then made in Sur-60 by systematically testing additional functional groups in the terminal naphthyl rings and the linker between the central phenyl rings. This optimization process resulted in substitution of one terminal naphthyl group with a phenalene group, addition of a hydroxyl group to the remaining sulfo-naphthyl group, and a carbonyl linker between the central rings to generate Sur-84 (−11.60 kcal/mol; **Fig. S6A**). Sur-84 makes additional contacts to His439 and Arg836 via one of the carboxylate groups and a hydrophobic interaction between the phenalene group and Val557. Finally, conversion of a central phenyl ring to naphthyl added hydrophobic contact with Tyr546 and generated Sur-85 (−11.96 kcal/mol; **Fig. 5C** and **S6A**).

The same process was also applied to the SARS-CoV-2 nucleocapsid RNA binding domain^29^ which has a shallow positively charged surface (**Fig. S5E-G**). Smaller suramin analogs with phenyl, stilbene and naphthyl moieties typically docked with higher scores than longer suramin analogs, with Sur-31 among the highest docking scores (−7.53 kcal/mol) and selected for further redesign to improve its affinity. Sur-31 is composed of two terminal naphthyl groups with a central unsubstituted phenyl ring linked connected by two conformationally restricted amide linkers (**Fig. S5H**). Inspection of the RNA binding pocket reveals Sur-31 is stabilized by multiple interactions: one sulfo-naphthyl ring with Arg93, Arg108 and Arg150, the central phenyl rings with Tyr110 and Tyr112, and the second sulfo-naphthyl ring with Tyr112, Thr92 and Arg89 (**Fig. 5D, E**). Our heparin screen also identified Hep-59 as binding to the same region, revealing potential for extending the Sur-31 scaffold at several sites (indicated * in **Fig. S5H**). Again, using docking-guided elaboration, one naphthyl ring was replaced by two sulfonyl-substituted phenyl rings connected to the central phenyl ring via a conformationally restricted azide linker resulting in Sur-114 with an improved of docking score of −8.79 kcal/mol (**Fig. S6B**). Sur-114 makes additional interactions with backbone carbonyls of Ala157 and Ile158, along with an increased van der Waals interaction with the protein. Further guided substitutions in the second naphthyl ring, adding another disulfonyl-hydroxy naphthyl ring linked by a flexible linker, generated Sur-121 with a further improvement in docking score to −9.30 kcal/mol (**Fig. S6B**). Final optimization of this naphthyl ring with a sulfate group resulted in Sur-127, with a final docking score of −9.34 kcal/mol. Sur-127 engages the overall same set of residues but with increased H-bonding and electrostatic interactions compared to Sur-31 (**Fig. 5F** and **S6B**).

This process thus identified an initial suramin analog for both SARS-CoV-2 protein targets, RdRp (Sur-36) and nucleocapsid (Sur-31), for which predicted affinity could be substantially improved by heparin docking-guided redesign efforts, resulting in final analogs Sur-85 and Sur-127, respectively. Overall, heparin docking- and structure-guided redesign of suramin analogs resulted in a collection of potential leads with markedly improved docking scores and other properties, compared to analogs identified in precision docking (compare green and purple scores, respectively in **Fig. 5G**).

We next investigated protein residue conservation to gain insights into the evolutionary constrains on the potential inhibitor binding sites in SARS-CoV-2 RdRp and nucleocapsid proteins. Across all coronaviruses the identified binding sites appear moderately and very highly conserved in nucleocapsid and RdRp, respectively (**Fig. 6**). As such, the phylogenetic relationship reveals the high similarity of both SARS-CoV-2 proteins to homologs in other β-coronavirus family members, comprising predominantly bat-origin viruses such as MERS-CoV and HCoV-HKU1 (**Fig. 6A, B; Fig S7**). The conservation of the RdRp binding site also extends to α-coronavirus family members of bat origin, which originated the human pathogens HCoV-229E and HCoV-NL63 (**Fig. 6A, B**). More broadly, the generally strong conservation in each binding site implies that the designed suramin analogs, Sur-85 and Sur-127, would potentially be equally effective in targeting these proteins in different coronaviruses. Thus, together the ligand docking/ elaboration strategies and evolutionary analyses provide a basis for designing specific inhibitors with broad anti-coronaviral activity.

**Figure 6.**
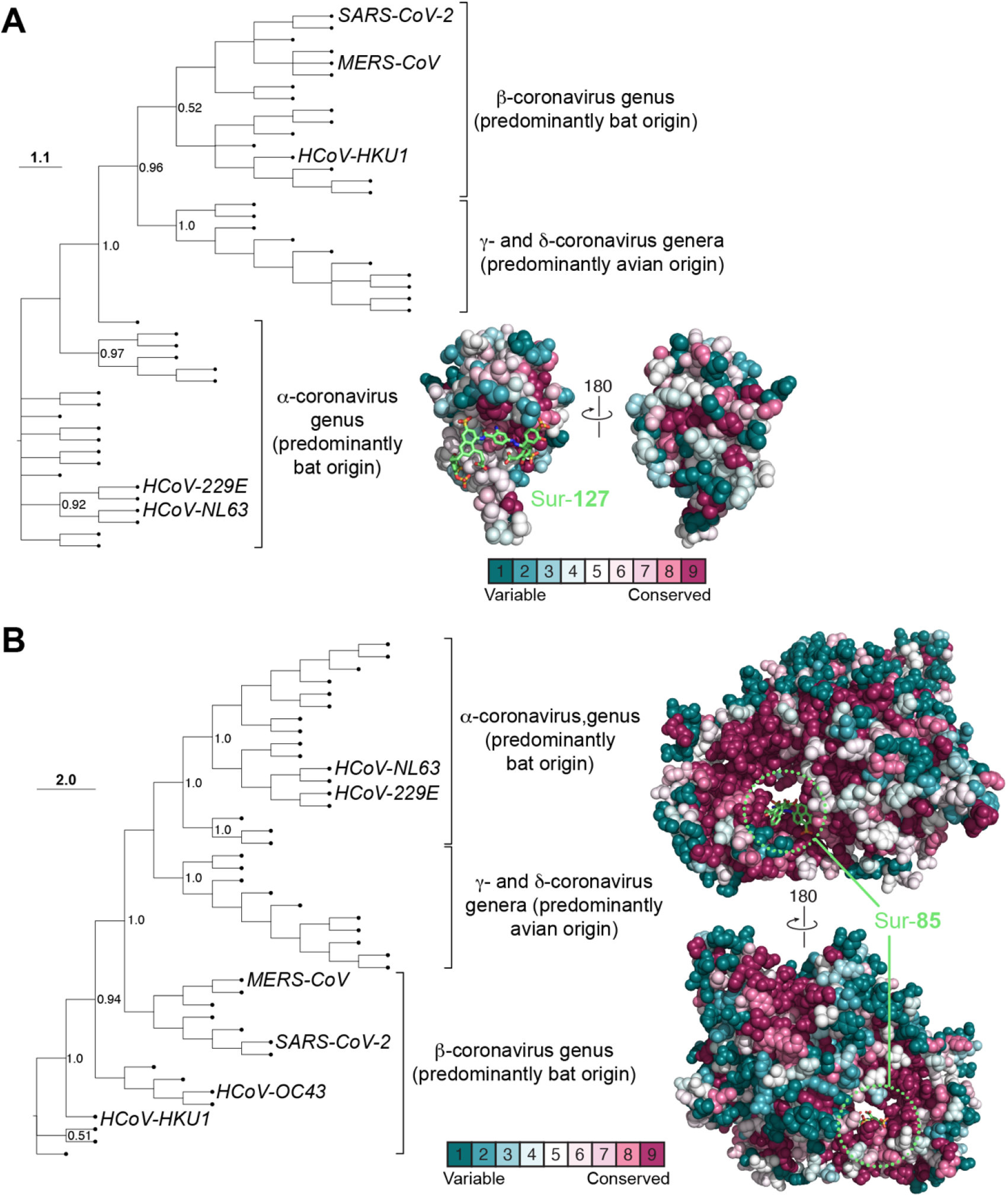
Maximum likelihood phylogenetic tree of coronavirus RdRp and nucleocapsid. ***A***, Phylogenetic tree of coronavirus capsid protein showing a separate clade for the β-coronavirus genus, with SARS-CoV-2 and MERS-CoV, which are mostly of bat origin. This clade is closely related to γ coronaviruses of avian origin while another bat origin α-coronavirus family forms a separate clade. Conservation of residues in coronavirus capsid proteins shows the binding pocket targeted by the designed suramin analog Sur-127 is evolutionarily conserved. ***B***, As *panel A* but for SARS-CoV-2 RdRp and Sur-85.

### Correlation of SP Glide docking and MM-GBSA rescoring for known suramin analogs

Computational pipelines for discovery of new compounds, such as the one described herein, rely on fast methods such as simple docking scoring functions for initial screening of large chemical libraries on a reasonable timescale. Among these approaches, the level of precision (e.g., HTVS vs SP Glide) also dictate the speed of docking calculations and thus our choice of initial HTVS followed by SP Glide rescoring (**Fig. 2** and **Fig. 3E,F**). However, even compound rankings from the more accurate SP Glide docking may fail to correlate with experimental activity or binding studies due to contributions of (unmodeled) waters, ions and other biological factors, as well as inherent flexibility of the protein target binding site in addition to ligand flexibility. More sophisticated approaches, such as free energy perturbation (FEP) or thermodynamic integration methods, can be applied at later stages of lead optimization, but are comparatively computationally expensive and thus not readily applicable for initial screening of large libraries. MM-GBSA is an alternate force field-based method which offers relatively quick binding free energy calculations^38^ and could be applied to rescoring ligand docking at one of several stages in our workflow.

To assess the utility of incorporating MM-GBSA calculations, we first used experimental data for suramin analogs targeted to SARS-CoV-2 RdRp, which were published during the course of this work^21^. First, SP Glide docking and MM-GBSA calculations were performed for suramin and five of the reported compounds^21^ and both methods showed good correlation with observed activity, particularly for MM-GBSA (R^2^ 0.63 for SP Glide and 0.90 for MM-GBSA; (**Fig. 7A, B**). The top published suramin analog for SARS-CoV-2 RdRp (NF157) with a reported IC_50_ of 0.041 μM yielded a MM-GBSA score of −69.7 kcal/mol. We next calculated MM-GBSA scores for the suramin analogs from our targeted re-design approach, and plotted these values to estimate their potential inhibitory activity. Many of our designed compounds show high MM-GBSA scores (< −50 kcal/mol) with potentially better than 0.3 μM predicted IC_50_. In particular, our re-design strategy based on Sur-36 (Sur-60, Sur-84 and Sur-85, shown as green points in **Fig. 7B**) result in a final compound with predicted IC_50_ that improves on the best currently reported and experimentally tested analog.

**Figure 7.**
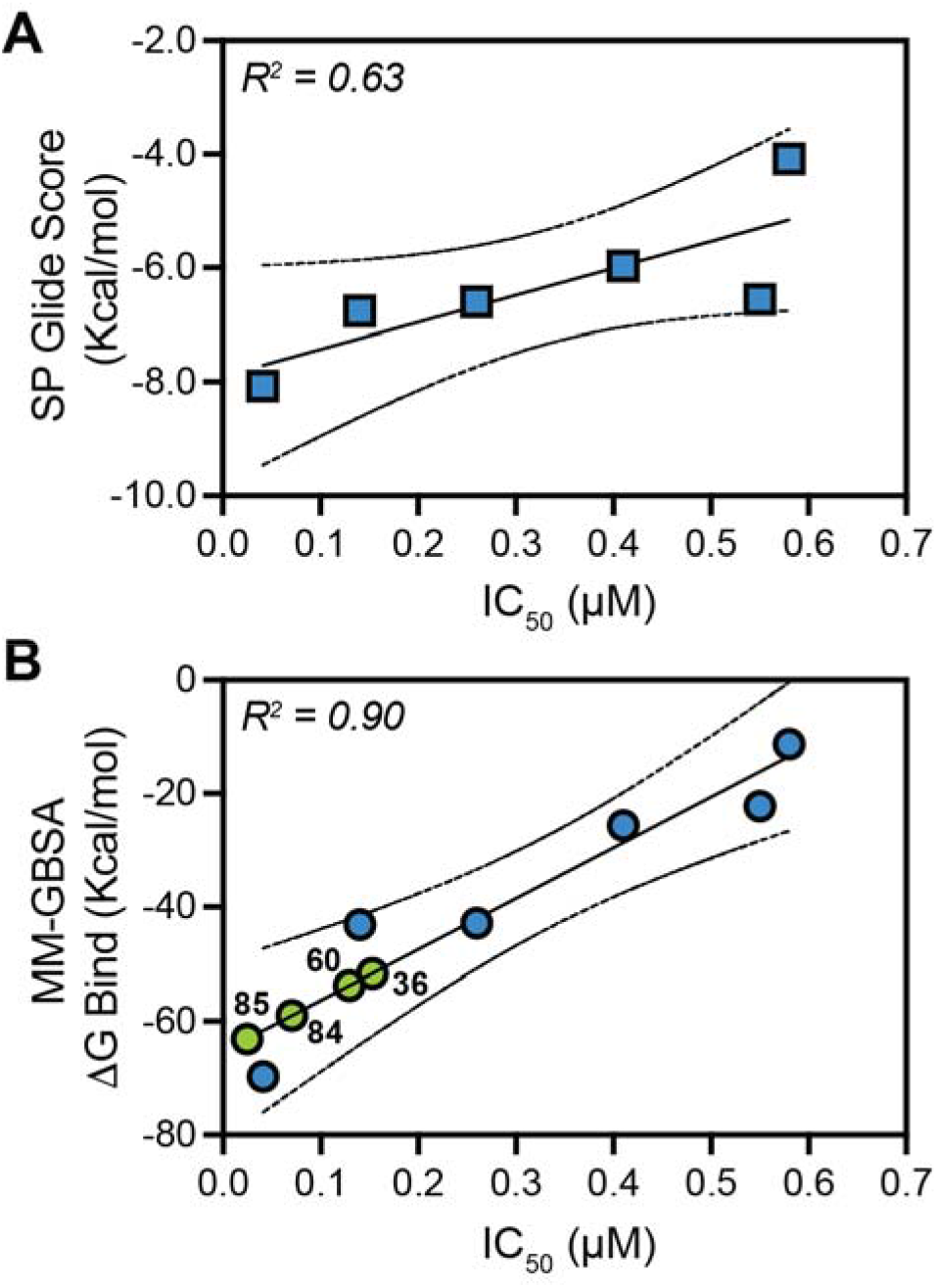
Correlation of suramin analog SP Glide docking and MM-GBSA rescoring for SARS-CoV-2 RdRp. Correlation between ***A***, SP Glide scores or ***B***, MM-GBSA with reported IC_50_ data^21^ for suramin and analogs with activity against SARS-CoV-2. The respective correlation coefficients (R^2^ values) for the linear fit to the data are shown on each plot. Values (labeled green points in *panel B*) for new suramin analogs designed based on Sur-36 (Sur-60, Sur-84 and Sur-85) are plotted using the linear fit to predict their potential activity.

Our additional protein targets for suramin analog re-design do not currently have experimental activity data for similar comparisons, so we therefore used three additional available datasets for suramin analogs with experimental binding or activity data for other protein targets. Suramin analogs have been reported that are capable of inhibiting the minichromosome maintenance protein 10 (Mcm10)^39^, an essential factor for DNA unwinding and a promising drug for anti-cancer therapy. Human Mcm10 was homology modeled based on *Xenopus* Mcm10 (PDB 3EBE) to allow SP Glide docking scores and MM-GBSA calculations for this system. Again, both approaches gave good correlation with experimentally determined binding affinities (K_d_ values), with SP-Glide performing marginally better (line fit R^2^ = 0.72 and 0.79 for MM-GBSA and SP Glide, respectively; **Fig. S8A**). Next, we used SIRT1, for which suramin analogs with inhibitory activity have been reported^23^, modeling the dimer interface using the SIRT5-suramin complex and SIRT1 crystal structures (PDB 2NYR and 4I5I, respectively). Our rescoring analysis reveals good correlation (R^2^ = 0.88) with MM-GBSA, but essentially no correlation with SP Glide score (**Fig. S8B**). Similarly, heparanase inhibitors which have potential use in inflammatory diseases, were reported based on suramin scaffold^40^. Our analysis (**Fig. S8C**) again showed no correlation between SP Glide and measured activity, whereas MM-GBSA was highly predictive (R^2^ = 0.92) of IC_50_, except for a small group of ligands (red points in **Fig. S8C**). We speculate that these outliers are a result of differential effect of those ligands on the target; for example, optimal binding of these ligands might exploit protein side chain flexibility of beyond the 5 Å radius of our calculations, or some other critical features such as hydration or bound ion.

## DISCUSSION

Suramin has garnered wide clinical interest for treating parasitic infections, as well as diseases ranging from viral infections (e.g. HIV, Dengue and Ebola) to human cancers^12^. Despite not satisfying typical drug likeliness criteria, such as Lipinski’s “rule of five”, suramin and its analogs have found success in clinical application. Suramin is negatively charged and binds predominantly positively charged surfaces in proteins involved in DNA and RNA processing, e.g., DNA and RNA polymerases, telomerase and chromodomain proteins, histone methyltransferases, and sirtuin histone deacetylases^19, 21, 23, 27^. Suramin also inhibits several membrane channels and signaling proteins. Despite suramin having been used for more than 100 years and shown to inhibit diverse protein families, there still exists a significant gap in our understanding of its promiscuous antagonist properties which likely also underpin its off-target and side effects in clinical use.

Prior to the current work, the nature of preferred binding surfaces and molecular determinants of suramin promiscuity were unclear. We reasoned that understanding these features could facilitate structure-based design of new suramin analogs with increased target specificity. Collectively, our analyses of available structures with suramin or suramin analogs revealed the contributions of both the physicochemical nature of the target protein binding pocket and conformational properties of suramin in making it a promiscuous ligand for diverse protein targets. In particular, suramin’s high flexibility, conferred by ten rotatable bonds, largely unsubstituted phenyl groups, and numerous non-directional electrostatic interactions with positively charged residues, collectively contribute to an ability to bind to structurally diverse nucleic acid interaction surfaces. Additionally, our analyses revealed an enrichment of aromatic residues in favored suramin binding sites and the role of various types of π-mediated interactions.

To date, suramin has been used as a lead in only a limited number of SAR studies. In one case, suramin-derived fragments were designed, synthesized and modified to act as an antagonist of the G protein-coupled receptor P_2_Y_2_ (ref.^41^), which mediates microglial inflammation. In another study, the toluene ring in suramin was substituted to improve suramin’s binding to falcipain-2 (ref.^42^), a cysteine protease of the malaria parasite *Plasmodium falciparum*. Although both these experimental studies used suramin as a lead, to our knowledge no high-throughput virtual screening has been performed nor a structure-guided approach taken to improve the affinity and selectivity of suramin based inhibitors. Here, we devised a virtual screening and precision docking strategy using sets of suramin and heparin analogs, the latter of which were then used to guide the redesign of top suramin analogs for enhanced target affinity and specificity. This redesign strategy was fully exemplified using SARS-CoV-2 nucleocapsid and RdRp proteins in a proof-of-concept redesign of suramin analogs as specific inhibitors of these viral proteins with potentially broad anti-coronaviral activity.

Suramin has been explored for its inhibitory activity against both bacterial and viral pathogens. Here, we demonstrated the potential utility of our virtual screening and ligand docking enabled identification and redesign of suramin analogs to improve their affinity and specificity for both known (RecA and HU) and new antibacterial targets (Fis and TsnR). For example, our study produced suramin analogs which bind more tightly to HU, based on computational docking scores, than those previously identified by screening inhibitors of HU^33^. Similarly, our computational strategy generated docking models for suramin and its analogs that can explain their previously identified capacity to block the function of diverse viral proteins including HIV-1 gp120, norovirus RdRp and bunyavirus nucleocapsid.

SARS-CoV-2 has also been shown to be inhibited by suramin^7^ and, during preparation of this manuscript, cryo-EM structural analysis and biochemical evidence revealed the molecular basis of suramin binding to SARS-CoV-2 RdRp^21^. Our docking study is consistent with these structural insights in which multiple binding poses of suramin were found in the RNA binding channel, one blocking the RNA template strand and the other binding to the primer strand in the catalytic site. Suramin and its analogs have been shown to be 20-fold more potent than remdesivir binding to the same site, and our results suggest that further improvement in suramin analog design is possible. Specifically, our docking-guided elaboration of suramin analogs showed improvement in the drug scaffold to make it a tighter binder based on both SP Glide docking and MM-GBSA rescoring calibrated using the experimentally determined binding data^21^. Using additional published datasets for suramin analogs with experimental binding or activity data for other protein targets, we also showed that MM-GBSA calculations consistently correlate well with experimental data. These studies thus suggest that incorporation of MM-GBSA at some stage of our computational pipeline (e.g., either prior to, or during lead analog redesign; **Fig. 2C**), can at a minimum enhance confidence in a ligand set identified by SP Glide, while for other targets its use may be essential. We speculate that the flexibility of the target, which is better modeled in MM-GBSA as compared to SP Glide scores, can be essential for success with some targets. However, initial selection of lead compounds from large libraries, can still reliably use much faster HTVS and SP Glide scoring, followed by MM-GBSA (or other more sophisticated approaches) to support later stages of lead optimization.

Our studies also identified the potential to target SARS-CoV-2 nucleocapsid with suramin analogs and proposed analogs possessing enhanced affinity and specificity. This approach could also be applied to the 3CL protease which was recently identified as another possible suramin binding target^43^. Our phylogenetic and evolutionary analyses also suggest that the suramin binding pockets are evolutionary conserved in related coronavirus RdRp and nucleocapsid proteins which could make these suramin analogs more broadly applicable therapeutics against this group of viral pathogens.

In summary, this study has defined the basis of suramin promiscuity and the nature of its preferred protein target binding sites and established a computational pipeline (**Fig. 2**) to rapidly identify new suramin analogs targeting diverse viral and bacterial targets for novel antimicrobial development. Suramin analogs screened in this study (e.g. Sur-8, Sur-23, Sur-25 and Sur-33) are built from modular aromatic fragments which can be decorated with different polar or charged substitutions and connected by various linkers (e.g. amide, sulfonamide and diazene; **Fig S9**). These properties can be readily exploited for future fragment-based designs of suramin-like inhibitors of viral and bacterial nucleic acid binding proteins. As such, our strategy offers the possibility for expedient design of novel suramin analogs with improved properties and efficacy for synthesis and experimental confirmation of their potential as novel lead antimicrobials.

## METHODS

### Virtual screening, docking and conformational analysis

Suramin and heparin analog sets were created using chemoinformatic guided searches in PubChem with parameters: Tanimoto similarity of >90%, <40 rotatable bonds, and molecular weight < 2 kD. The structures of compounds retrieved were processed with the LigPrep module in Schrödinger Software. For conformational analysis of the ligands, the ConfGen module ^44^ of Schrödinger Software was used with the “mix torsional low mode sampling” method, which is a combination of *Monte Carlo* and systematic sampling of rotatable bonds to find low energy ligand conformations.

To prepare each protein (“receptor”) for ligand docking, hydrogen atoms were added, and the structure energy minimized using the Protein Preparation Wizard in Schrödinger Software. The docking grid location was determined by analysis of the nucleic acid binding region of each protein, either by prediction using SiteMap (Schrödinger Software) or from published experimental evidence for a given target protein. Docking grid size was determined based on the nucleic acid binding pocket volume and designed to completely cover all residues and centered symmetrically. Docking was performed using the virtual screening workflow of Glide^45^ in Schrödinger Software, in HTVS mode and then for precision docking in SP mode as used in our previous studies^33, 46^. MM-GBSA calculations were performed in the Prime module of Schrödinger Software with the OPLS3e force field and allowing protein side chain flexibility within a 5 Å radius of the bound ligand.

Ligand physicochemical properties and chemical similarity were calculated in the Canvas module of Schrödinger Software ^47^. Ligand-protein nonbonded interactions were calculated using BIOVIA DS Viewer for salt bridges, hydrogen bonds and other π mediated interactions. PyMOL was used for the visual inspection of structures and generation of figures.

### Phylogenetic analysis and conservation study

Homologous sequences of coronavirus RdRp and nucleocapsid were collected by BLAST search in NCBI with ~10,000 and 5,000 sequences identified, respectively. The large number of sequences arises due to the number of sequenced SARS-CoV-2 and related viruses that have >99% identity in the dataset. Therefore, the CD-HIT server^48^ was next used to remove redundant sequences from the dataset using a sequence identity cut-off of 98%. Multiple sequence alignment and evolutionary analyses were then performed using Clustal Omega and MEGA 7 (ref.^49^), respectively, with evolutionary history inferred using the Maximum Likelihood method with bootstrap consensus tree inferred from 100 replicates. The fraction of replicate trees in which the associated taxa clustered together in the bootstrap test (100 replicates) are shown next to the branches. Evolutionary distances were computed using the JTT matrix-based method and are in the units of number of amino acid substitutions per site. The rate variation among sites was modeled with a gamma distribution (shape parameter 4). Finally, the phylogenetic tree was visualized using FigTree (http://tree.bio.ed.ac.uk/software/figtree/). Conserved residues were plotted on the SARS-CoV-2 RdRp (PDB 7BV1, chain A) and nucleocapsid (PDB 6M3M, chain A) using ConSurf^50^ from the pre-calculated multiple sequence alignment.

## Supporting information

Supplementary File-Top 100 LIgans SP Glide Docking

## ASSOCIATED CONTENT

### Supplementary Results

Precision docking of suramin and heparin analogs to bacterial and viral protein targets.

### Supplementary Figures

**Figure S1**. Structural basis of viral protein inhibition by suramin.

**Figure S2.** Docking-guided elaboration of top suramin analogs against TsnR and Fis.

**Figure S3.** Docking-guided elaboration of top suramin analogs against RecA and HIV gp120.

**Figure S4.** Docking-guided elaboration of top suramin analogs against norovirus RdRp and bunyavirus nucleocapsid.

**Figure S5.** Docking-guided elaboration of top suramin analogs against SARS-CoV2 RdRp and nucleocapsid.

**Figure S6.** Suramin analog redesign using iterative chemical enumeration and docking.

**Figure S7.** Phylogenetic analysis of coronavirus RdRp and nucleocapsid.

**Figure S8.** Correlation of SP Glide docking and MM-GBSA rescoring for known suramin analogs

**Figure S9.** Building blocks of suramin analogs.

### Supplementary Tables

**Table S1.** Protein crystal structures with bound suramin and suramin analogs.

**Table S2.** Ligands used as database search query for suramin and heparin analogs.

**Table S3.** Glide SP docking score of suramin analogs to bacterial and viral targets.

**Table S4.** Glide SP docking score of heparin analogs to bacterial and viral targets.

**Table S5.** Glide SP docking score of computationally designed suramin analogs based on Sur-31 and Sur-37 against SARS-CoV-2 RdRp and nucleocapsid

### Additional Supplementary Files

Dey-JCIM-Supplementary File-Top 100 Ligands SP Glide Docking.xlsx

Dey-JCIM-Supplementary File-SMILES.csv

Atomic coordinates of protein targets with suramin or heparin analogs from ligand docking studies. Files are named by target and ligand using the naming convention described in the main text, e.g. *HU-Sur1.pdb*.

### Data and Software Availability

Additional underlying data from these studies beyond those provided in the Supplementary Materials will be made available on request by the authors.

## AUTHOR INFORMATION

### Author Contributions

DD, SR and GLC conceptualized the project. DD and GLC wrote the draft. All authors have given approval to the final version of the manuscript.

## ACKNOWLEDGEMENT

We thank Dr. William Wuest for comments on the manuscript and members of the Conn lab for discussions and comment through the course of this work.

