## Supporting Information for "Targeted redesign of suramin analogs for novel antimicrobial lead development"

**Precision docking of suramin and heparin analogs to bacterial and viral protein targets.**

***Thiostrepton-resistance Methyltransferase (TsnR)*.** TsnR (PDB 3GYQ) possesses a large positively charged cleft (**Fig. S2**), structurally similar to HU, for interaction with the bacterial 23S ribosomal RNA. TsnR is bound with highest affinity by Sur-5 (docking score -9.95 kcal/mol), a tyrosyl-phenylalanine peptidomimetic ligand. However, this ligand was not selected for further optimization due to its high number of rotatable bonds and thus inherent flexibility and greater potential for promiscuity. Sur-6 binds (docking score -9.86 kcal/mol) to the positively charged cleft of TsnR in an extended conformation (**Fig. S2C**), similar that observed for Sur-1 bound to HU (**Fig. 4A**). Sur-6 has a double sulfophenyl core with terminal dihydroxybenzoic acid terminal rings connected by conformationally rigid diazene linkers (**Fig. S2D**). The overall symmetric binding of the molecule to the positively charged RNA binding cleft, conformational restriction, and low overall charge makes Sur-6 a promising lead for TsnR inhibitor design. Superposition of the docking pose of heparin analog Hep-42 (docking score -12.12 kcal/mol) in the same binding pocket identifies multiple positions (marked * in **Fig. S2D**) suitable for targeted resign of Sur-6 to further improve affinity and target selectivity.

***DNA-binding protein Fis (Fis)*.** Fis (PDB 3JR9) is functionally similar to HU and also engages DNA via a positively charged cleft. Among the top scoring suramin analogs, Sur-13 was selected for further analysis as it adopts the most symmetrical binding pose in the target pocket (**Fig. S2E**-**G**). Hep-45 also binds to Fis (docking score -10.52 kcal/mol) in a similar pose as Sur-13, again identifying multiple potential locations for modification Sur-13 with additional substituents to increase its affinity and specificity for the Fis binding pocket (marked * in **Fig. S2H**).

***DNA recombination protein RecA (RecA)*.** RecA (PDB 4OQF) is a monomeric DNA-binding protein with a smaller but deeper and more asymmetric binding interface as compared to the other target proteins^1^ (**Fig. S3**). The docking analyses show that RecA is bound by suramin and its very close analog Sur-15 with these ligands in a strongly bent conformation (shown for Sur-15 in **Fig. S3C**). This bent conformation is facilitated by the rotatable bonds between the terminal and central rings in one half of the ligand. Heparin analog Hep-50 also binds RecA (docking score -9.26 kcal/mol) with a similar binding pose to Sur-15. Substitution of the central phenyl rings of Sur-15 (marked * in **Fig. S3D**) could potentially increase its affinity and engage residues in the pocket which were identified from interactions made with Hep-50.

***Human immunodeficiency virus glycoprotein 120 (HIV gp120)*.** HIV gp120 (6IEQ) binds to smaller compounds than those identified for the other targets. HIV gp120 binds to asymmetric and conformationally restricted suramin analog Sur-22 with high score (docking score -6.10 kcal/mol; **Fig. S3E-G**). Sur-22 is composed of a sulfo-naphthyl ring in the center, with substituted benzenesulfonic acids in the terminal region (**Fig. S3H**). The docking pose with hyaluronic acid analog Hep-51 overlaps partially with Sur-22 and, once again, reveals potential sites for ligand redesign (marked * in **Fig. S3H**) with a sulfonyl phenyl ring or other polar substituent that could increase the affinity.

***Norovirus RNA-dependent RNA polymerase (RdRp) and bunyavirus nucleocapsid*.** Suramin and its analogs bind to norovirus RdRp (**Fig. S4A, B**) and bunyavirus nucleocapsid (**Fig. S4E, F**) and disrupt the viral replication and host attachment processes^2, 3^. Our post-docking analysis of Norovirus RdRp (PDB 3UR0) shows that suramin analog Sur-24 binds to the RNA binding site in a bent conformation (**Fig. S4C**). Our docking study also shows that the same analog can bind in different conformations to the RdRp channel. This flexibility of suramin and its analogs in the norovirus RdRp channel could be the reason that only partial electron density for suramin was observed in norovirus RdRp bound structure (**Fig. S1A**). Hep-56 binding to RdRp has partial overlap but reveals a different pose than the suramin analogs (**Fig. S4C**). The RdRp binding site is large and different classes of molecule can thus take different poses to block the RNA binding channel. Nonetheless, overlay of the Sur-34 and Hep-56 docking poses identified positions in both the central and terminal rings of Sur-34 (**Fig. S4D**) which could be substituted with polar extensions to increase its specificity and affinity. A similar analysis of the bunyavirus nucleocapsid identified Sur-27 as binding with highest score (docking score -9.48 Kcal/mol; **Fig. S4G,H**); this analog is similar to suramin with a sulfonyl amide linker. Heparin analogs with sulfonyl groups bind more tightly to nucleocapsid protein with Hep-52 showing the highest score of -10.79 kcal/mol. Our virtual screening and precision docking thus provides a model of interaction of suramin and its analogs with RdRp and nucleocapsid and also identifies new analogs and a rational path for redesign for specific targets.

**FIGURES**

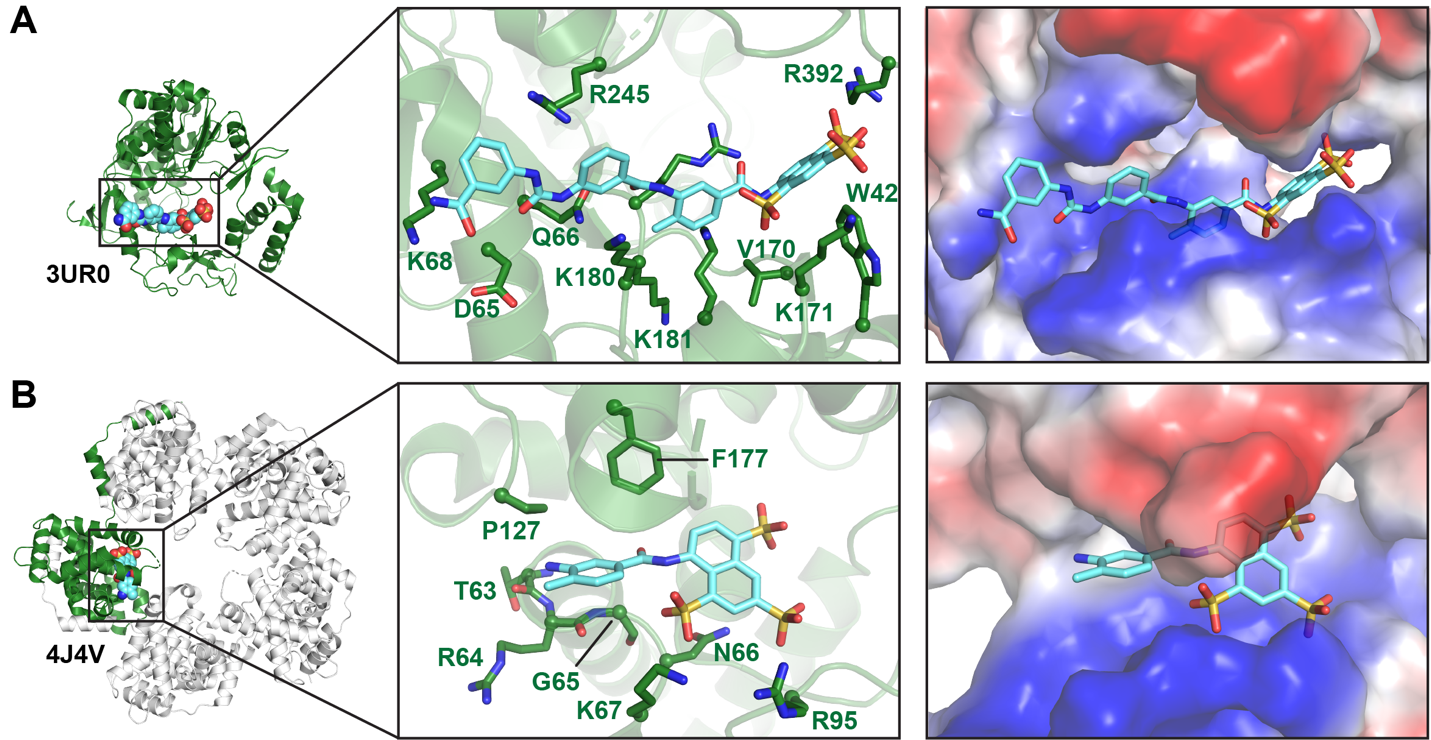

**Figure S1. Structural basis of viral protein inhibition by suramin. *A***, modeled suramin fragment in complex with norovirus RdRp (PDB 3UR0) showing its interaction with a predominantly positively charged surface (electrostatic surface potential is shown in the right image) including residues K68, K171, K180, K181 and R392 in the RNA binding cleft. ***B****,* As *panel A* but for bunyavirus nucleocapsid protein (PDB 4J4V) and its RNA binding region containing positively charged residues K67 and R95.

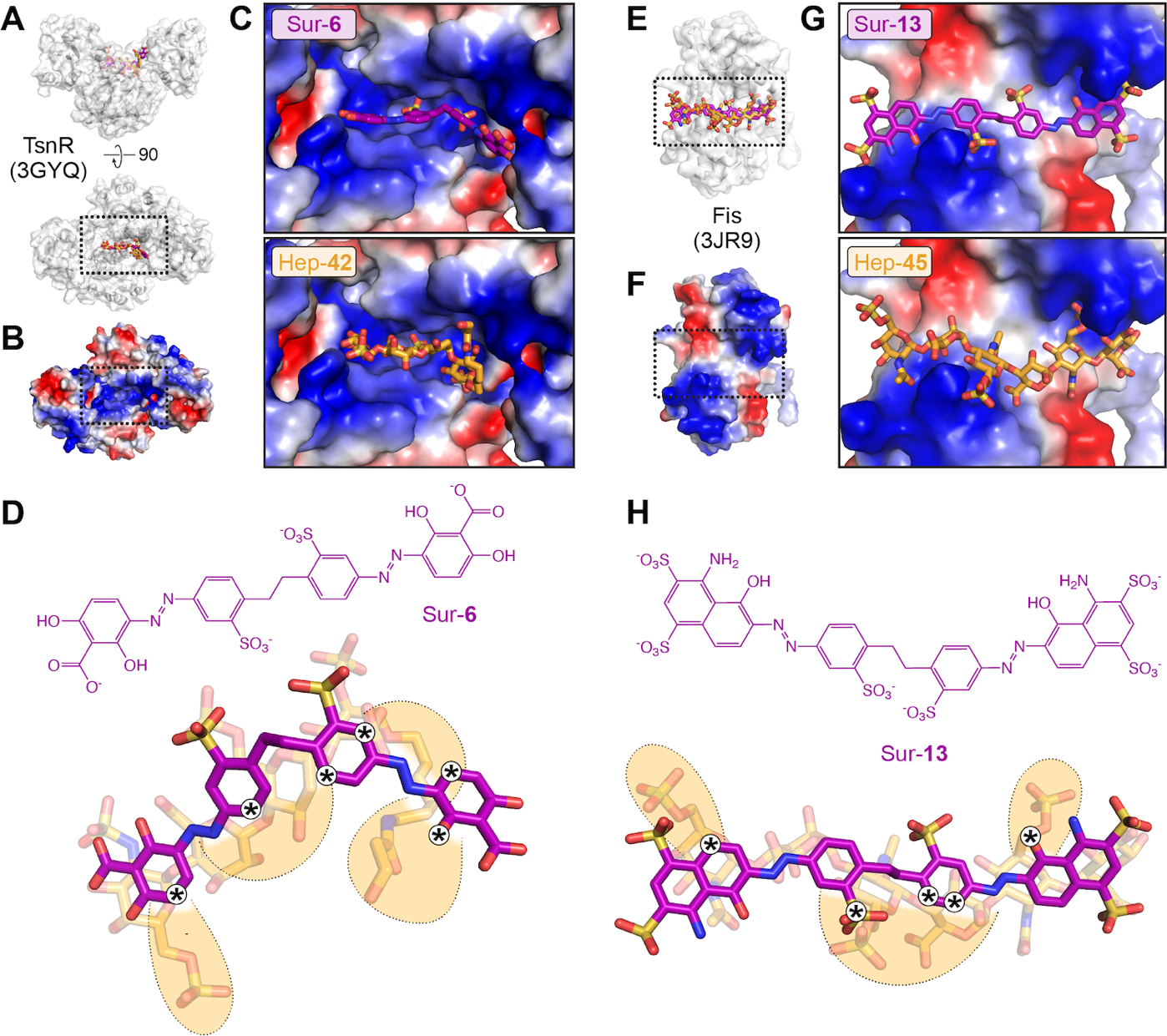

**Figure S2. Docking-guided elaboration of top suramin analogs against TsnR and Fis. *A***, Overview of SP Glide docking of suramin analog Sur-6 (purple sticks) and Hep-42 (orange sticks) in the TsnR dimer interface (white surface). ***B***, TsnR dimer shown with electrostatic surface potential. ***C*,** Zoomed in views of the boxed region of *panels A* and *B* with Sur-6 and Hep-42 bound at overlapping sites in the positively charged interface of the TsnR dimer. ***D***, *Top*, chemical structure of Sur-6. *Bottom*, the overlapping binding pose for Hep-42 used to guide identification of sites (marked *) in Sur-6 for potential redesign to improve binding affinity and specificity. ***E****-****H***, as for *panels A-D* but for Sur-13 and using Hep-45 to guide its redesign to target Fis.

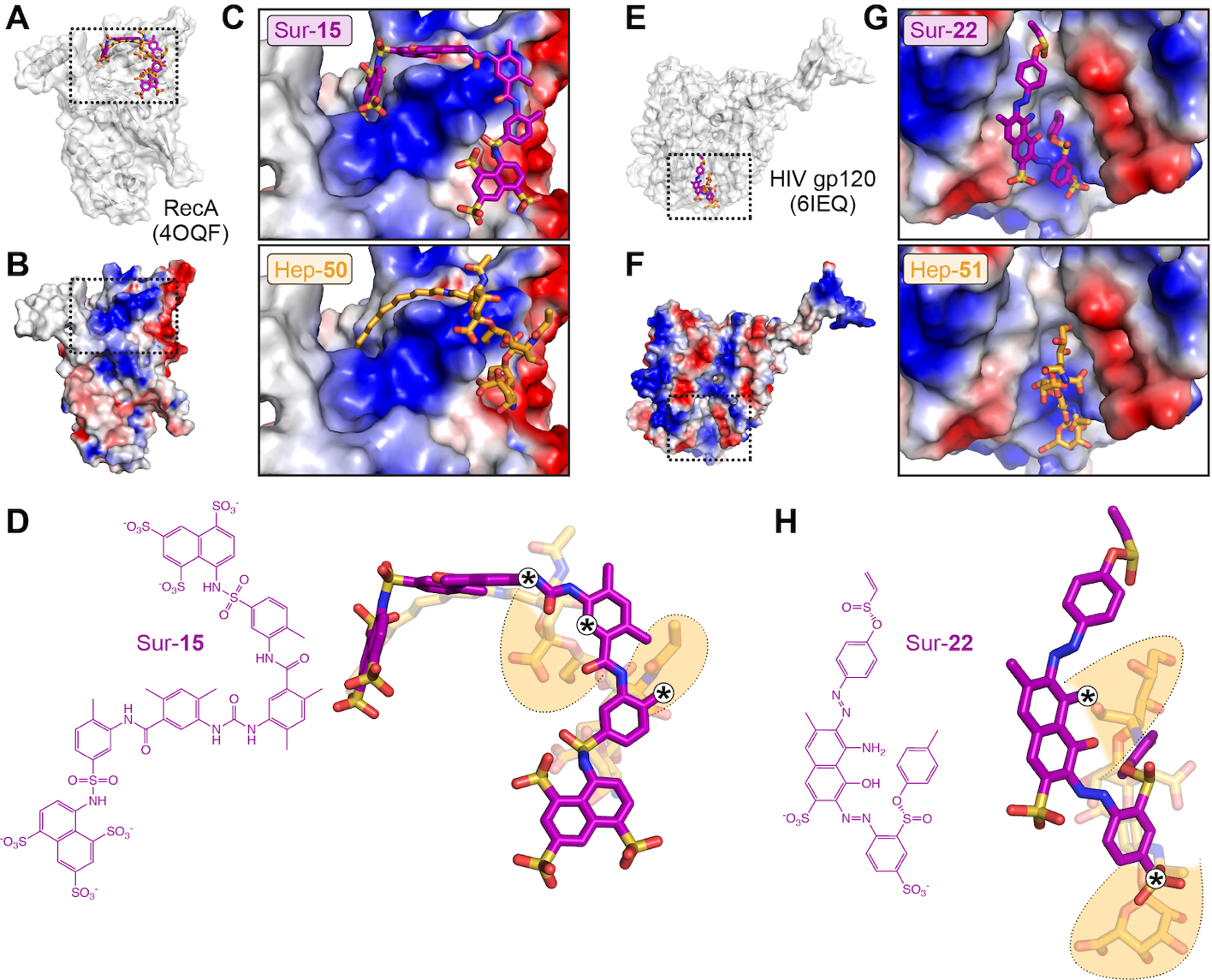

**Figure S3. Docking-guided elaboration of top suramin analogs against RecA and HIV gp120. *A***, Overview of SP Glide docking of suramin analog Sur-15 (purple sticks) and Hep-50 (orange sticks) with RecA (white surface). ***B***, RecA shown with electrostatic surface potential. ***C*,** Zoomed in views of the boxed region of *panels A* and *B* with Sur-15 and Hep-50 bound at overlapping sites at a positively charged surface of RecA. ***D***, *Top*, chemical structure of Sur-15. *Bottom*, the overlapping binding pose for Hep-50 used to guide identification of sites (marked *) in Sur-15 for potential redesign to improve binding affinity and specificity. ***E****-****H***, as for *panels A-D* but for Sur-22 and using Hep-51 to guide its redesign to target HIV gp120.

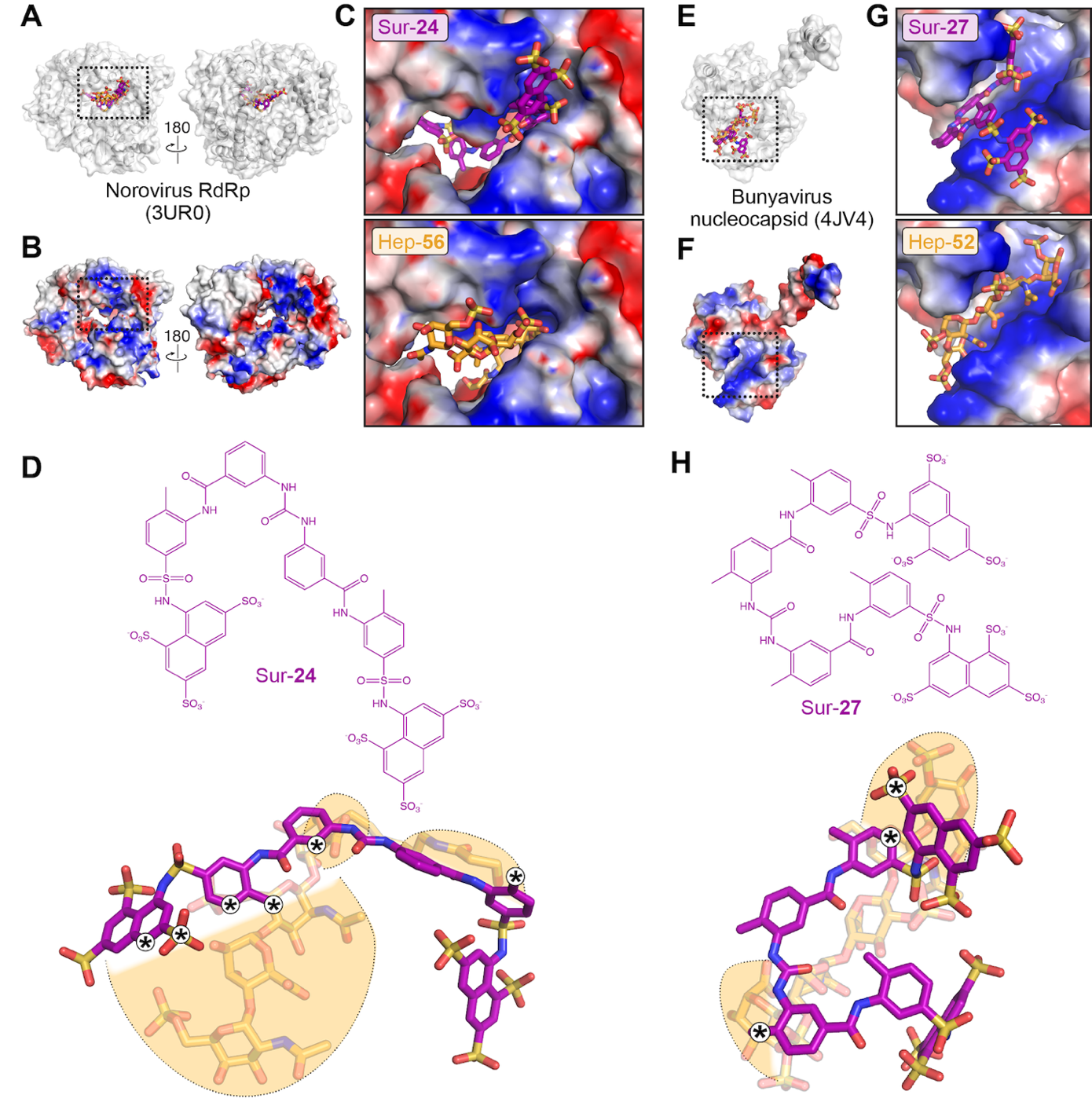

**Figure S4. Docking-guided elaboration of top suramin analogs against norovirus RdRp and bunyavirus nucleocapsid. *A***, Overview of SP Glide docking of suramin analog Sur-24 (purple sticks) and Hep-56 (orange sticks) with norovirus RdRp (white surface). ***B*,** Norovirus RdRp shown with electrostatic surface potential. ***C***, Zoomed in views of the boxed region of *panels A* and *B* with Sur-24 and Hep-56 bound at overlapping sites on a positively charged surface of norovirus RdRp. ***D***, *Top*, chemical structure of Sur-24. *Bottom*, the overlapping binding pose for Hep-56 used to guide identification of sites (marked *) in Sur-24 for potential redesign to improve binding affinity and specificity. ***E****-****H***, as for *panels A-D* but for Sur-27 and using Hep-52 to guide its redesign to target bunyavirus nucleocapsid.

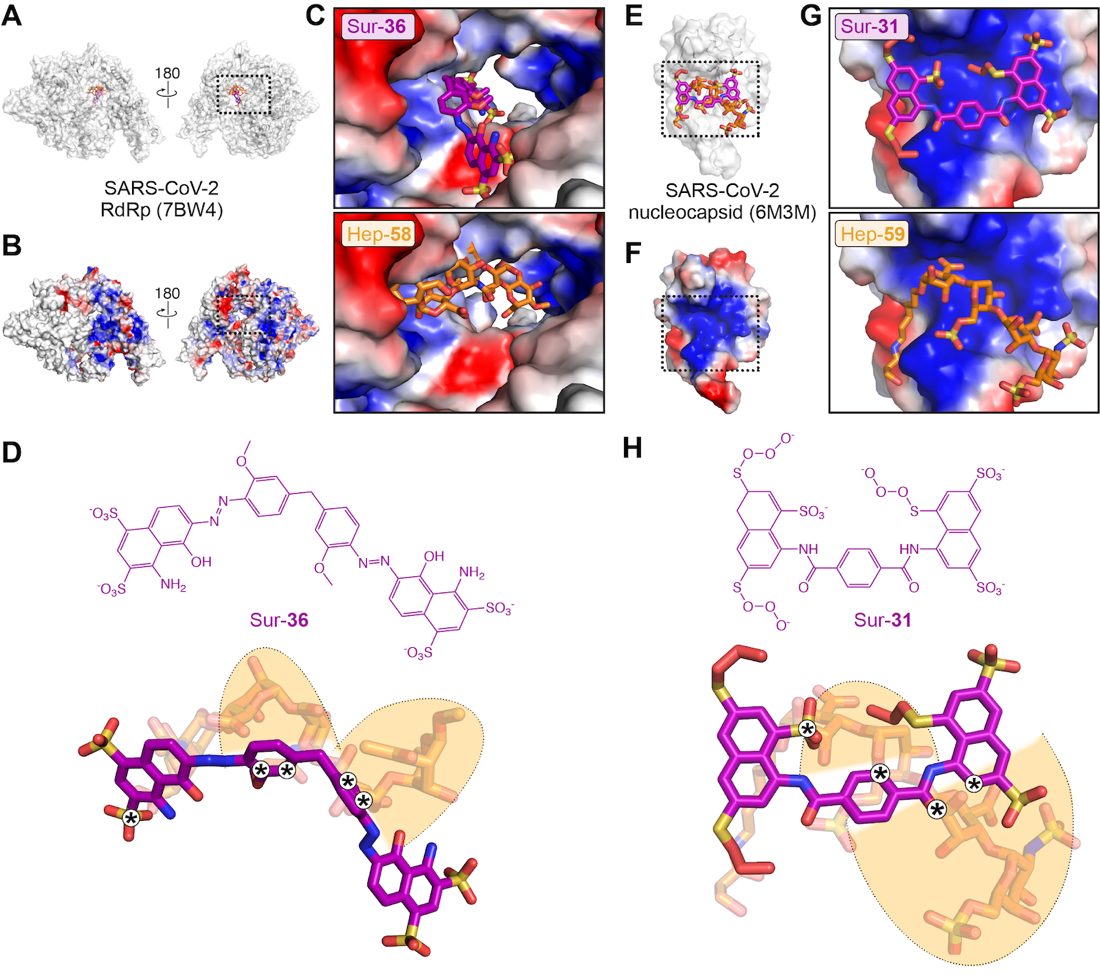

**Figure S5. Docking-guided elaboration of top suramin analogs against SARS-CoV2 RdRp and nucleocapsid. *A***, Overview of SP Glide docking of suramin analog Sur-36 (purple sticks) and Hep-58 (orange sticks) with SARS-CoV-2 RdRp (white surface). ***B***, SARS-CoV-2 RdRp shown with electrostatic surface potential. ***C***, Zoomed in views of the boxed region of *panels A* and *B* with Sur-36 and Hep-58 bound at overlapping sites on a positively charged surface of SARS-CoV-2 RdRp. ***D***, *Top*, chemical structure of Sur-36. *Bottom*, the overlapping binding pose for Hep-58 used to guide identification of sites (marked *) in Sur-36 for potential redesign to improve binding affinity and specificity. ***E****-****H***, as for *panels A-D* but for Sur-31 and using Hep-59 to guide its redesign to target SARS-CoV-2 nucleocapsid.

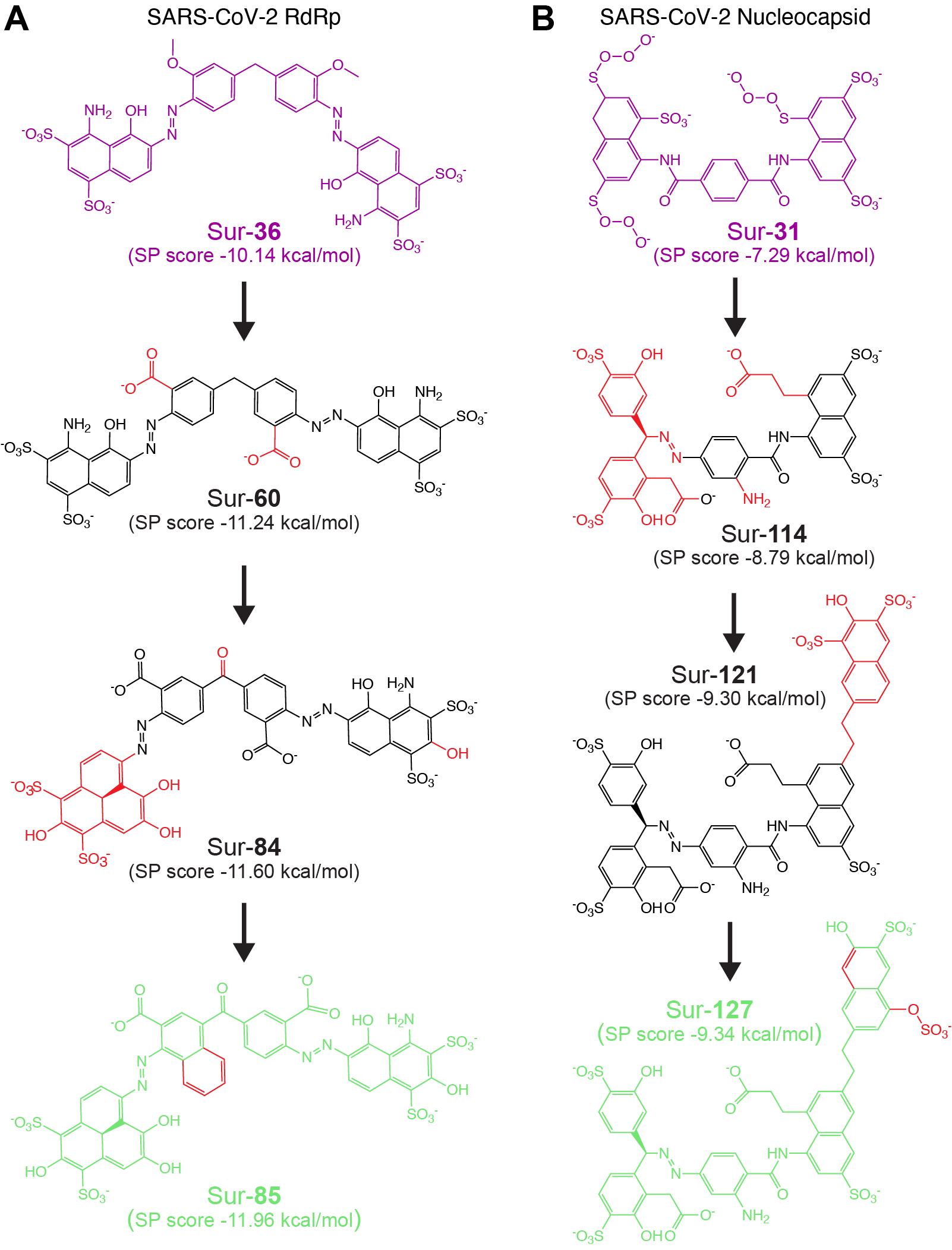

**Figure S6. Suramin analog redesign using iterative chemical enumeration and docking. *A***, Modification of Sur-36, the top scoring suramin analog against SARS-CoV-2 RdRp, by replacement of the methoxy groups of the central phenyl rings with carboxylate groups (to generate Sur-60), replacement of the terminal naphthyl group with a phenalene group and other modifications highlighted in red (Sur-84), and a further modification of the central ring (highlighted in red) (Sur-85). Each modification resulted in improved SP Glide score as compared to parent compound suramin and each prior analog. ***B***, The same strategy applied to targeting SARS-CoV-2 nucleocapsid, beginning from Sur-31. Replacement of one naphthyl ring (*left*) with two phenyl rings substituted with sulfonyl and hydroxyl groups connected to the central phenyl ring with conformationally restricted azide linker produced Sur-114); adding another disulfonyl-hydroxy naphthyl ring via a flexible linker to the second naphthyl right (*right*) generated Sur-121); and, final optimization resulted in Sur-127.

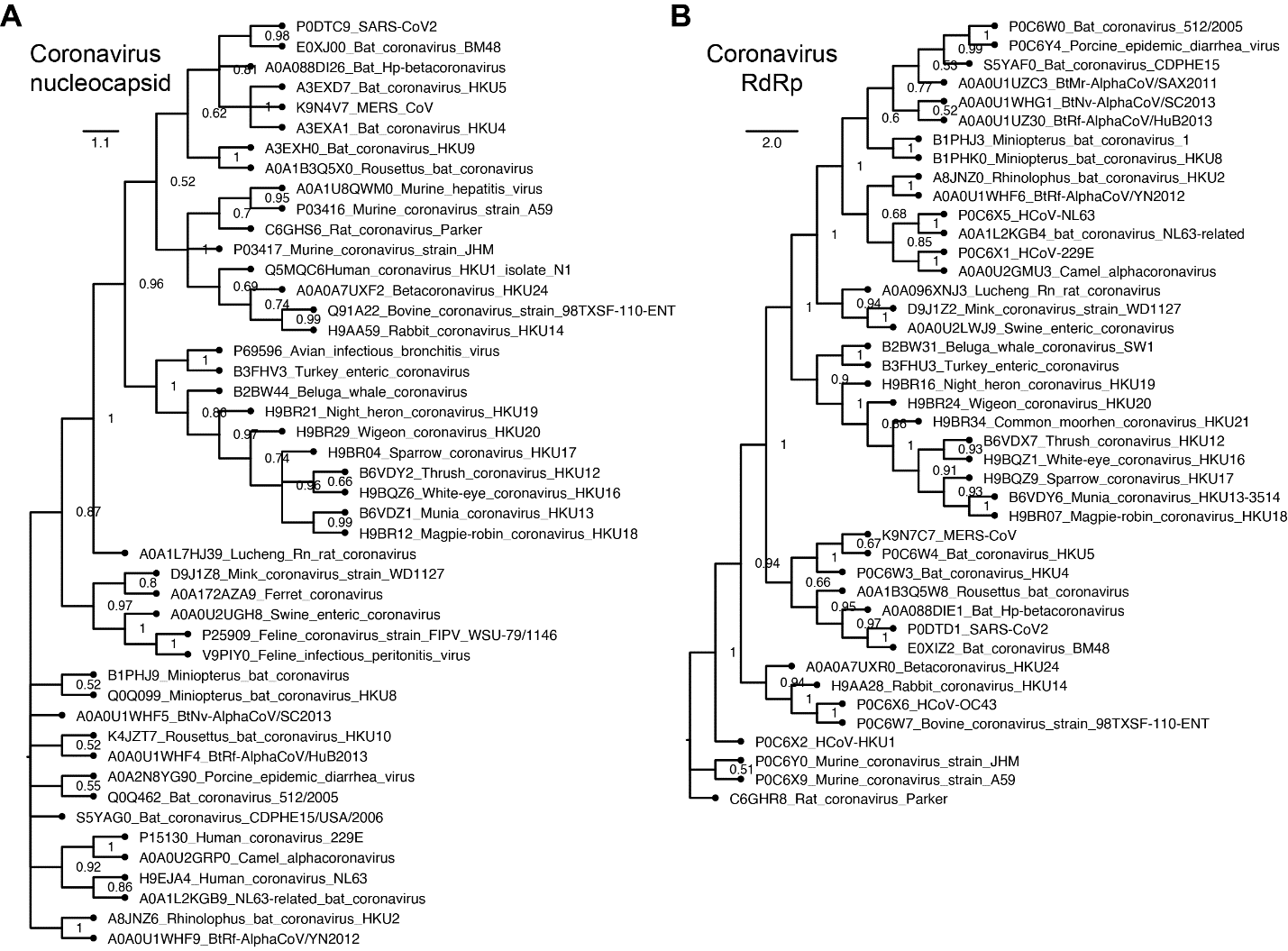

**Figure S7.** **Phylogenetic analysis of coronavirus RdRp and nucleocapsid.** Maximum likelihood phylogenetic tree for ***A***, coronavirus RdRp and ***B***, nucleocapsid protein. The edge shows bootstrap values where values closer to 1 indicate highest branch confidence. The scale bar for each tree shows average number of protein substitutions per site.

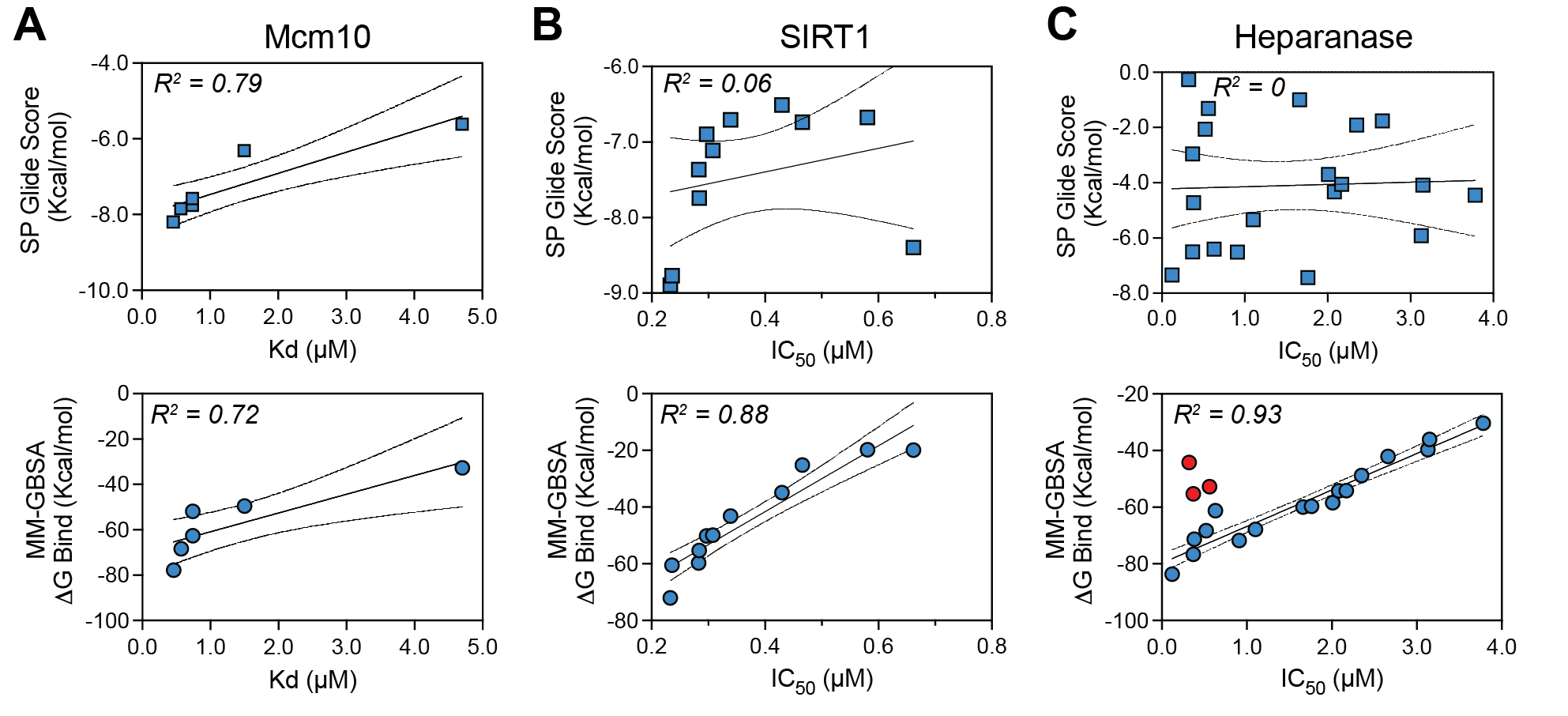

**Figure S8. Correlation of SP Glide docking and MM-GBSA re-scoring for known suramin analogs.** Correlation with experimental binding or activity data for ***A***, human Mcm10, ***B***, SIRT1 and ***C***, heparanase shown with the respective correlation coefficients (R^2^ values) on each plot. For Mcm10, both MM-GBSA (bottom row) and SP Glide (top row) show good correlation, whereas for SIRT1 and heparanase only MM-GBSA scoring resulted a good correlation with experimentally determined activities.

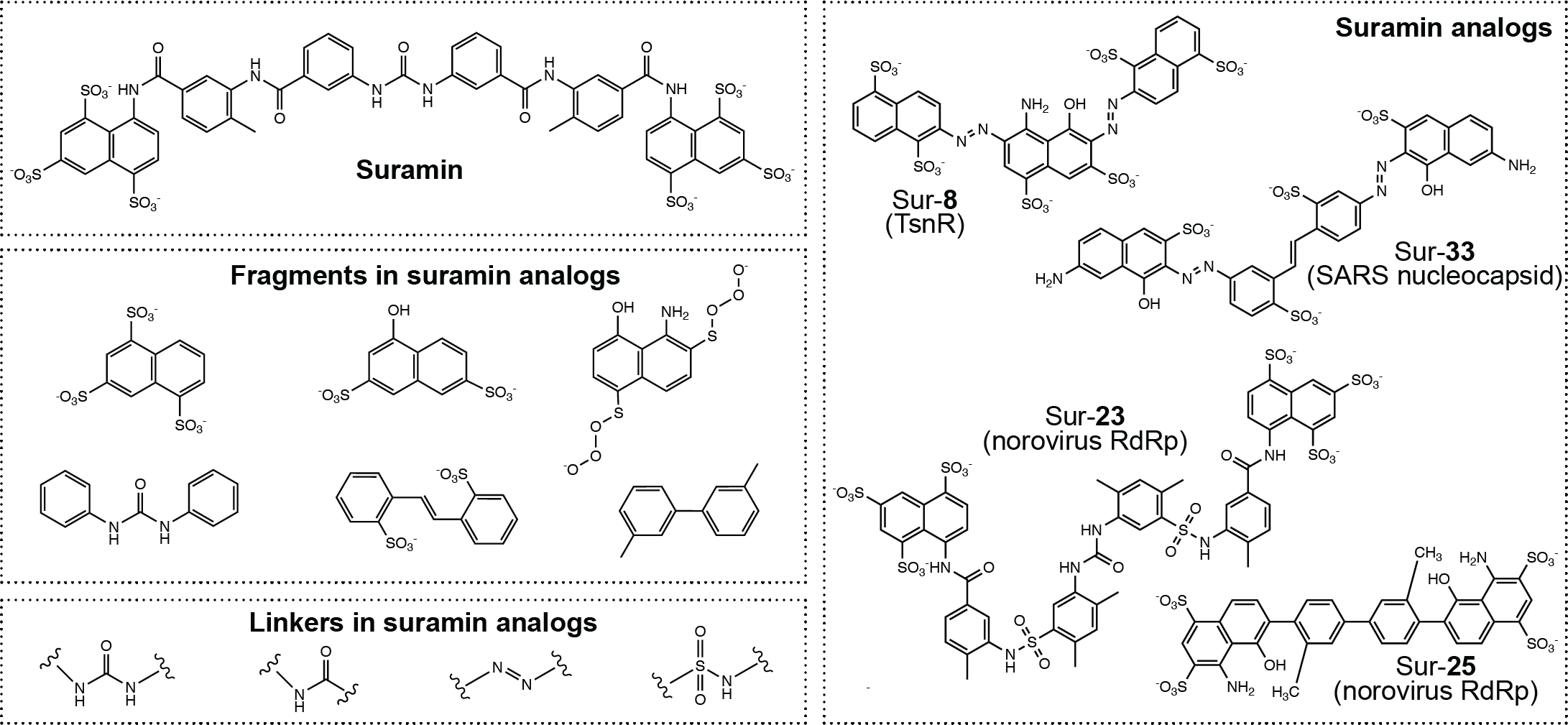

**Figure S9. Building blocks of suramin analogs.** Comparison of suramin with fragments and linkers found in suramin analogs. Different structural arrangements and combinations offer a wide scope for new analog design as shown for the examples (*right*) of top scoring analogs against different viral and bacterial protein targets.

| **Table S1.** **Protein crystal structures with bound suramin and suramin analogs** | | | |
| --- | --- | --- | --- |
| **Compound**  **(PubChem CID)** | **PDB** | **Interacting Protein** | **Organism** |
| Suramin (5361) | 3PP7 | Pyruvate kinase | *Leishmania mexicana* |
|  | 6CE2 | Myotoxin I (Phospholipase) | *Bothrops moojeni (snake)* |
|  | 1Y4L | Myotoxin II (Phospholipase) | *Bothrops asper* |
|  | 2NYR | NAD+-dependent deacetylase SIRT5 | *Human* |
|  | 3UR0 | RNA-dependent RNA polymerase (RdRp) | *Norovirus* |
|  | 4X3U | CBX7 chromodomain | *Human* |
|  | 3BF6 | Thrombin | *Human* |
|  | 4J4R | Nucleocapsid protein | *Phlebovirus JS2010-018 (Bunyavirus)* |
|  | 4J4V | Nucleocapsid protein | *Phlebovirus JS2010-018 (Bunyavirus)* |
| Ponceau S (80367) | 3QV9 | Pyruvate kinase | *Leishmania mexicana* |
| Acid blue 80 (20551) | 3QV6 | Pyruvate kinase | *Leishmania mexicana* |
| NF023 (6093160) | 3UPF | RNA-dependent RNA polymerase (RdRp) | *Norovirus* |
| Suramin analog (78673842) | 4NRT | RNA-dependent RNA polymerase (RdRp) | *Norovirus* |
| Suramin analog (78673842) | 4NRU | RNA-dependent RNA polymerase (RdRp) | *Norovirus* |

| **Table S2. Ligands used as database search query for suramin and heparin analogs** | | |
| --- | --- | --- |
| **Compound** | **PubChem CID** | **Scaffold/repeating unit** |
| Suramin | 5361 | Naphthalene-1,3,5-trisulfonic Acid |
| Y-ART-4 | 16130554 | Tyrosine, N'-,O'-decasulfate |
| Sb-1 | 370101 | Naphthalene-1,3,5-trisulfonic Acid |
| Evans Blue | 3315 | Naphthalene-1,3,5-trisulfonic Acid |
| NF449 | 6093161 | Naphthalene-1,3,5-trisulfonic Acid |
| Heparin | 772 | Sulfonylgalactose |
| Chondroitin sulfate | 24766 | Sulfonylgalactose |
| Beta-D-galactan | 53356679 | Galactose |
| Dextran sulfate | 3084656 | Dextran Trisulfate |

| **Table S3. Glide SP docking score of suramin analogs to bacterial and viral targets** | | | |
| --- | --- | --- | --- |
| **Suramin Analog #** | **PubChem CID** | **Docking score** | **Target Protein** |
| **Sur-1** | 136610971 | -9.67 | HU |
| **Sur-2** | 189950 | -9.27 | HU |
| **Sur-3** | 138395884 | -9.08 | HU |
| **Sur-4** | 54484026 | -9.07 | HU |
| **Sur-5** | 101450366 | -9.95 | TsnR |
| **Sur-6** | 135499186 | -9.86 | TsnR |
| **Sur-7** | 23332466 | -9.70 | TsnR |
| **Sur-8** | 136641721 | -9.38 | TsnR |
| **Sur-9** | 137285174 | -9.21 | TsnR |
| **Sur-10** | 136187549 | -9.90 | Fis |
| **Sur-11** | 54282040 | -9.27 | Fis |
| **Sur-12** | 135788852 | -8.99 | Fis |
| **Sur-13** | 366386 | -8.92 | Fis |
| **Sur-14** | 23332450 | -8.36 | Fis |
| **Sur-15** | 54460062 | -9.53 | RecA |
| **Sur-16** | 53946823 | -9.23 | RecA |
| **Sur-17** | 60161100 | -9.21 | RecA |
| **Sur-18** | 88775689 | -8.28 | RecA |
| **Sur-19** | 135462996 | -7.64 | RecA |
| **Sur-20** | 20046446 | -9.70 | HIV-gp120 |
| **Sur-21** | 136650873 | -6.28 | HIV-gp120 |
| **Sur-22** | 136144399 | -6.10 | HIV-gp120 |
| **Sur-23** | 53870817 | -11.00 | Norovirus RdRp |
| **Sur-24** | 6093160 | -10.51 | Norovirus RdRp |
| **Sur-25** | 10010648 | -10.29 | Norovirus RdRp |
| **Sur-26** | 136967510 | -9.86 | Norovirus RdRp |
| **Sur-27** | 54077393 | -9.49 | Bunyavirus Capsid |
| **Sur-28** | 86576527 | -9.35 | Bunyavirus Capsid |
| **Sur-29** | 5361 | -9.27 | Bunyavirus Capsid |
| **Sur-30** | 16760580 | -8.88 | Bunyavirus Capsid |
| **Sur-31** | 21053626 | -7.53 | SARS-CoV-2 Capsid |
| **Sur-32** | 136165952 | -7.28 | SARS-CoV-2 Capsid |
| **Sur-33** | 136644835 | -6.67 | SARS-CoV-2 Capsid |
| **Sur-34** | 136612800 | -6.65 | SARS-CoV-2 Capsid |
| **Sur-35** | 135962521 | -6.56 | SARS-CoV-2 Capsid |
| **Sur-36** | 137262334 | -10.14 | SARS-CoV-2 RdRp |
| **Sur-37** | 53875413 | -9.95 | SARS-CoV-2 RdRp |
| **Sur-38** | 54225859 | -9.63 | SARS-CoV-2 RdRp |

| **Table S4. Glide SP docking score of heparin analogs to bacterial and viral targets** | | | |
| --- | --- | --- | --- |
| **Heparin Analog #** | **PubChem CID** | **Docking score** | **Target Protein** |
| **Hep-39** | 91845952 | -10.02 | HU |
| **Hep-40** | 129176649 | -9.68 | HU |
| **Hep-41** | 772 | -9.53 | HU |
| **Hep-42** | 118439248 | -12.13 | TsnR |
| **Hep-43** | 57984120 | -11.17 | TsnR |
| **Hep-44** | 10844145 | -11.12 | TsnR |
| **Hep-45** | 42609008 | -10.54 | FIS |
| **Hep-46** | 71586046 | -10.38 | FIS |
| **Hep-47** | 57984119 | -10.49 | FIS |
| **Hep-48** | 138154530 | -9.91 | RecA |
| **Hep-49** | 138305172 | -9.47 | RecA |
| **Hep-50** | 88571766 | -9.26 | RecA |
| **Hep-51** | 130198279 | -8.36 | HIV-gp120 |
| **Hep-52** | 91862102 | -10.79 | Bunyavirus Capsid |
| **Hep-53** | 58106098 | -9.79 | Bunyavirus Capsid |
| **Hep-54** | 145769611 | -9.64 | Bunyavirus Capsid |
| **Hep-55** | 118439251 | -10.55 | Norovirus RdRp |
| **Hep-56** | 118439241 | -10.28 | Norovirus RdRp |
| **Hep-57** | 59228874 | -10.69 | SARS-CoV-2 RdRp |
| **Hep-58** | 59164169 | -10.00 | SARS-CoV-2 RdRp |
| **Hep-59** | 118439245 | -9.49 | SARS-CoV-2 Capsid |

| **Table S5. Glide SP docking score of computationally designed suramin analogs based on Sur-31 and Sur-37 against SARS-CoV-2 RdRp and nucleocapsid** | | |
| --- | --- | --- |
| **Suramin Analog #** | **Docking score** | **Target Protein** |
| Sur-60 | -11.25 | SARS-CoV-2 RdRp |
| Sur-61 | -10.33 | SARS-CoV-2 RdRp |
| Sur-62 | -10.26 | SARS-CoV-2 RdRp |
| Sur-63 | -10.97 | SARS-CoV-2 RdRp |
| Sur-64 | -10.14 | SARS-CoV-2 RdRp |
| Sur-65 | -10.40 | SARS-CoV-2 RdRp |
| Sur-66 | -11.49 | SARS-CoV-2 RdRp |
| Sur-67 | -10.10 | SARS-CoV-2 RdRp |
| Sur-68 | -10.83 | SARS-CoV-2 RdRp |
| Sur-69 | -10.42 | SARS-CoV-2 RdRp |
| Sur-70 | -10.36 | SARS-CoV-2 RdRp |
| Sur-71 | -11.67 | SARS-CoV-2 RdRp |
| Sur-72 | -10.12 | SARS-CoV-2 RdRp |
| Sur-73 | -10.36 | SARS-CoV-2 RdRp |
| Sur-74 | -11.28 | SARS-CoV-2 RdRp |
| Sur-75 | -10.53 | SARS-CoV-2 RdRp |
| Sur-76 | -11.39 | SARS-CoV-2 RdRp |
| Sur-77 | -11.31 | SARS-CoV-2 RdRp |
| Sur-78 | -10.94 | SARS-CoV-2 RdRp |
| Sur-79 | -10.86 | SARS-CoV-2 RdRp |
| Sur-80 | -10.58 | SARS-CoV-2 RdRp |
| Sur-81 | -10.14 | SARS-CoV-2 RdRp |
| Sur-82 | -10.01 | SARS-CoV-2 RdRp |
| Sur-83 | -11.36 | SARS-CoV-2 RdRp |
| Sur-84 | -11.60 | SARS-CoV-2 RdRp |
| Sur-85 | -11.96 | SARS-CoV-2 RdRp |
| Sur-86 | -10.62 | SARS-CoV-2 RdRp |
| Sur-87 | -11.72 | SARS-CoV-2 RdRp |
| Sur-88 | -10.51 | SARS-CoV-2 RdRp |
| Sur-89 | -11.57 | SARS-CoV-2 RdRp |
| Sur-90 | -10.86 | SARS-CoV-2 RdRp |
| Sur-91 | -11.15 | SARS-CoV-2 RdRp |
| Sur-92 | -11.13 | SARS-CoV-2 RdRp |
| Sur-93 | -10.92 | SARS-CoV-2 RdRp |
| Sur-94 | -10.67 | SARS-CoV-2 RdRp |
| Sur-95 | -10.60 | SARS-CoV-2 RdRp |
| Sur-96 | -10.57 | SARS-CoV-2 RdRp |
| Sur-97 | -10.52 | SARS-CoV-2 RdRp |
| Sur-98 | -10.10 | SARS-CoV-2 RdRp |
| Sur-99 | -11.12 | SARS-CoV-2 RdRp |
| Sur-100 | -7.32 | SARS-CoV-2 Capsid |
| Sur-101 | -7.11 | SARS-CoV-2 Capsid |
| Sur-102 | 7.02 | SARS-CoV-2 Capsid |
| Sur-103 | -8.29 | SARS-CoV-2 Capsid |
| Sur-104 | -8.03 | SARS-CoV-2 Capsid |
| Sur-105 | -7.78 | SARS-CoV-2 Capsid |
| Sur-106 | -7.65 | SARS-CoV-2 Capsid |
| Sur-107 | -7.65 | SARS-CoV-2 Capsid |
| Sur-108 | -7.46 | SARS-CoV-2 Capsid |
| Sur-109 | -5.88 | SARS-CoV-2 Capsid |
| Sur-110 | -5.54 | SARS-CoV-2 Capsid |
| Sur-111 | -5.48 | SARS-CoV-2 Capsid |
| Sur-112 | -5.18 | SARS-CoV-2 Capsid |
| Sur-113 | -5.13 | SARS-CoV-2 Capsid |
| Sur-114 | -8.79 | SARS-CoV-2 Capsid |
| Sur-115 | -8.26 | SARS-CoV-2 Capsid |
| Sur-116 | -8.20 | SARS-CoV-2 Capsid |
| Sur-117 | -8.05 | SARS-CoV-2 Capsid |
| Sur-118 | -7.92 | SARS-CoV-2 Capsid |
| Sur-119 | -7.74 | SARS-CoV-2 Capsid |
| Sur-120 | -7.45 | SARS-CoV-2 Capsid |
| Sur-121 | -9.30 | SARS-CoV-2 Capsid |
| Sur-122 | -8.79 | SARS-CoV-2 Capsid |
| Sur-123 | -8.44 | SARS-CoV-2 Capsid |
| Sur-124 | -7.95 | SARS-CoV-2 Capsid |
| Sur-125 | -7.20 | SARS-CoV-2 Capsid |
| Sur-126 | -6.42 | SARS-CoV-2 Capsid |
| Sur-127 | -9.33 | SARS-CoV-2 Capsid |
| Sur-128 | -8.62 | SARS-CoV-2 Capsid |
| Sur-129 | -8.24 | SARS-CoV-2 Capsid |
| Sur-130 | -9.18 | SARS-CoV-2 Capsid |
| Sur-131 | -9.15 | SARS-CoV-2 Capsid |
| Sur-132 | -9.12 | SARS-CoV-2 Capsid |
| Sur-133 | -8.60 | SARS-CoV-2 Capsid |
